# Multi-Step Pathway Engineering in Probiotic *Saccharomyces boulardii* for Abscisic Acid Production in the Gut

**DOI:** 10.1101/2024.12.22.629964

**Authors:** Femke Van Gaever, Paul Vandecruys, Yasmine Driege, Seo Woo Kim, Johan M. Thevelein, Rudi Beyaert, Jens Staal

**Author notes:** These authors share last authorship (Rudi Beyaert, Jens Staal). **Author Contributions** F.V.G., P.V., J.M.T., R.B., and J.S. conceived the experiments. F.V.G., Y.D., S.W.K. and J.S. performed the experiments. F.V.G. analyzed the data. F.V.G., P.V., J.M.T., R.B. and J.S. discussed the results. F.V.G. and J.S. wrote the manuscript. F.V.G., P.V., J.M.T., R.B. and J.S. revised the manuscript.

## Abstract

The plant hormone abscisic acid (ABA) has gained attention for its role in animals and humans, particularly due to its protective effects in various immune and inflammatory disorders. Given its high concentrations in fruits like figs, bilberries and apricots, ABA shows promise as a nutraceutical. However scalability, short half-life and cost limit the use of ABA-enriched fruit extracts and synthetic supplements. In this study, we propose an alternative ABA administration method to overcome these challenges. We genetically engineered a strain of the probiotic *Saccharomyces boulardii* to produce and deliver ABA directly to the gut of mice. Using the biosynthesis pathway from *Botrytis cinerea*, four genes (*bcaba1-4*) were integrated into *S. boulardii*, enabling ABA production at 30°C, as previously described in *Saccharomyces cerevisiae*. Introducing an additional cytochrome P450 reductase gene resulted in a 7-fold increase in ABA titers, surpassing previous ABA-producing *S. cerevisiae* strains. Supplementation of the ABA-producing *S. boulardii* in the diet of mice (at a concentration of 5 x 10^8^ CFU/g) led to effective gut colonization but resulted in low serum ABA levels (approximately 1.8 ng/mL). The absence of detectable serum ABA after administration of the ABA-producing probiotic through oral gavage, prompted further investigation to determine the underlying cause. The physiological body temperature (37°C) was identified as a major bottleneck for ABA production. Modifications to enhance the mevalonate pathway flux improved ABA levels at 37°C. However, additional modifications are needed to optimize ABA production before testing this probiotic in disease contexts in mice.

## Background

Abscisic acid (ABA) is a well-characterized phytohormone involved in the regulation of key physiological functions, including responses to abiotic stress and plant growth and development (1). Although the effects of ABA have been extensively studied within the plant kingdom, its role in animals has only gained significant attention over the past decades (2). In animals, the involvement of ABA has been reported in various immune and inflammatory responses (3), and treatments of animals and humans with ABA have demonstrated protective effects against several diseases, such as colitis (4), type 2 diabetes (5,6), atherosclerosis (7) and depression (8). Given its high concentrations in fruits like figs, bilberries and apricots (6), ABA presents potential as an interesting nutraceutical substance (2). However, challenges such as scalability, short half-life and cost-effectiveness hinder the practical use of ABA-enriched fruit extracts and synthetic ABA supplements (9,10). In this study, we propose an alternative ABA administration technique that we believe could effectively address these challenges.

Probiotics are defined as “microbial cell preparations or components of microbial cells that have a beneficial effect on the health and well-being of the host” (11). Leveraging decades of advancements in metabolic engineering, next-generation engineered probiotics can offer unique attributes for delivering biomolecules to the gut (12–14). Engineered probiotics have traditionally been bacterial, with *Escherichia coli* Nissle 1917 modified to eliminate pathogens like *Pseudomonas aeruginosa* (15) or to treat conditions such as hyperammonemia (16) and cancer (17). However, a major disadvantage of bacterial probiotics is their susceptibility to antibiotics and bacteriophages (18). In addition to bacteria, the gastrointestinal tract also contains a diverse fungal population (19). In this study, we focus on the non-pathogenic yeast *Saccharomyces boulardii*, a variety of the budding yeast *Saccharomyces cerevisiae*. *S. boulardii* is widely recognized as safe and commonly used as a commercial probiotic for treating diarrhea (20) and various gut-related diseases (21–27), with its probiotic effects largely attributed to its unusually high acetic acid production (28). Through genetic engineering, *S. cerevisiae* has been modified to produce various plant-based products (9), including artemisinic acid, a precursor of the antimalarial drug artemisinin (29,30). As well-established protocols developed for *S. cerevisiae* can be applied to *S. boulardii* (31–33), this enables efficient genetic modification. Previous genetic engineering efforts have enabled *S. boulardii* to secrete lysozyme (33) and HIV-1 gag (34) in cell culture, the vitamin A precursor β-carotene in the gut of germ-free mice (35), and IL-10 or elevated levels of acetic acid to exert protection in mouse models of inflammatory bowel disease (36–38).

In plants, ABA is synthesized via the degradation of C40 carotenoids produced in plastids (39). However, plants are not the only organisms capable of producing ABA. ABA production has also been confirmed in phytopathogenic fungi such as *Botrytis cinerea* (40), cyanobacteria (41), the animal parasite *Toxoplasma gondii* (42) and mammals, including humans (43). Although ABA signaling properties are conserved across different kingdoms of life, biosynthetic pathways vary among organisms. The only well-understood non-plant ABA synthesis pathway has been described in the grey mold *B. cinerea*, where ABA is synthesized from the C15 molecule farnesyl pyrophosphate (FPP), a product from the mevalonate (MVA) pathway (44,45).

Specifically, FPP is cyclized to form the ABA core scaffold and subsequently oxidized by multiple enzymes (45,46). In 2006, the gene cluster involved in these subsequent steps was identified to contain two cytochrome P450 monooxygenase (CYP) genes (*bcaba1* and *bcaba2*), a short-chain dehydrogenase/reductase gene (*bcaba4*), and an alpha-ionylideneethane synthase gene (*bcaba3*) (47–49). Recently, Otto *et al.* demonstrated that integrating the ABA biosynthesis gene cluster from *B. cinerea* (47) into *S. cerevisiae* (50) did not encounter limitations in precursor supply for ABA production. Instead, the bottlenecks in this pathway were identified as the activities of CYPs BcABA1 and BcABA2. Further overexpression of the cytochrome P450 reductase (CPR) gene *bccpr1*, which facilitates electron transfer from NADPH to CYPs (51), increased ABA titers (50). The probiotic properties of *S. boulardii*, combined with the recent report demonstrating that *S. cerevisiae* can be genetically engineered to produce ABA (50), suggest that engineering *S. boulardii* to produce ABA could offer an efficient strategy for ABA delivery to the gut.

## Results

### Establishment of an ABA-producing *S. boulardii* strain

In this study, our objective was to engineer a highly active and fast-growing genetically modified probiotic *S. boulardii* strain, with minimal modifications while achieving maximal ABA production levels. We selected two specific *S. boulardii* strains for this purpose: the Enterol strain, commercialized by Biocodex and commonly used in therapeutic applications, and the high acetic acid-producing SbP strain, known for its potential as a probiotic candidate due to the antimicrobial effects of acetic acid (28,52). Based on the findings of Otto *et al.,* which indicated that the introduction of *Botrytis cinerea* genes *bcaba1*, *bcaba2*, *bcaba3* and *bcaba4* enables ABA production in *S. cerevisiae*, we integrated these four genes into specific predefined integration sites (53) in the genomes of the *S. boulardii* strains (28) (**Table S1**). This genetic modification resulted in the creation of the E-ABA1 and SbP-ABA1 strains, respectively (**Table 1**). Following 24 h of cultivation at 30°C in 50 mL of yeast extract peptone dextrose (YPD) medium, supernatants of the wild-type strains and the engineered strains were collected. A reporter assay optimized by our group (54) was used to measure ABA concentrations in culture supernatants, with (+)-ABA used to generate a standard curve. An ABA production of 8.5 mg/L and 8.6 mg/L was measured in culture supernatant of the E-ABA1 and SbP-ABA1 strains, respectively (**Figure 1A**). These measurements confirmed similar levels of ABA production in both the Enterol and SbP strains of *S. boulardii* expressing *bcaba1-4*. Notably, a decrease in maximum optical density (OD_600_) was observed between the ABA-producing strains and the background strains, suggesting a potential trade-off between growth and ABA production (**Figure S1A**).

**Figure 1.**
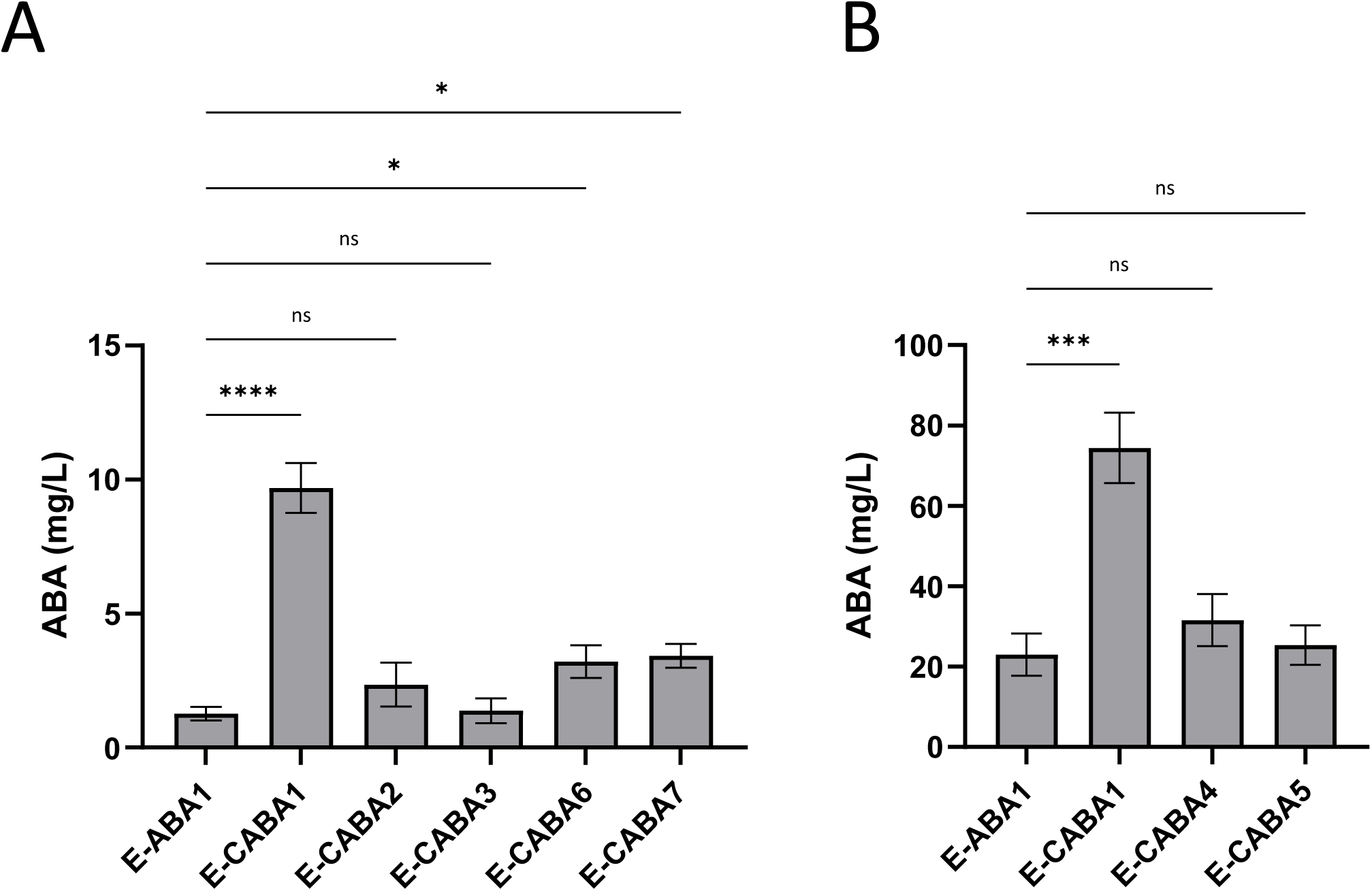
ABA production in *S. boulardii* and *S. cerevisiae* strains *in vitro* at 30°C. **(A)** ABA titer in supernatant of ABA-producing strains. E-ABA1 is based on the genetic background of *S. boulardii* strain Enterol (E) and contains *bcaba1-4* and SbP-ABA is based on the genetic background of high acetic acid-producing *S. boulardii* strain SbP (SbP). Strains were cultivated for 24 h at 30°C in 50 mL of YPD medium. Average ABA titers were calculated from 5 independent biological replicates. **(B)** ABA titer in supernatant of strains E-ABA1 and TABA3. E-ABA1 is based on the genetic background of *S. boulardii* strain Enterol and contains *bcaba1-4*, TABA3 (50) is based on *S. cerevisiae* strain CENK.PK113-5D and contains *bcaba1-4*, *bccpr1* and *tHMG1*. Strains were cultivated for 12 h or 24 h at 30°C in 20 mL of YPD medium. Average ABA titers were calculated from 3 independent biological replicates. Data are represented as mean ± SEM and statistically analyzed using a Student’s t-test. * p < 0.05, ** p < 0.01, *** p < 0.001, **** p < 0.0001.

**Table 1.** Strains used in this study.

|  | Strain name | Genotype | Source / Reference |
| --- | --- | --- | --- |
| Background strains | SbP | Wild type *S. boulardii* strains | Lene Jespersen, University of Copenhagen, Denmark (1) |
|  | Enterol (E) | Wild type *S. boulardii* isolated from Pharmacy product Enterol® | Johan Thevelein, KU Leuven, Belgium (2) |
| Strains with *B. cinerea* genes | TABA3 | *S. cerevisiae* CEN.PK113-5D<br>*P<sub>PGK1</sub>.bcaba1 P<sub>PGK1</sub>.bcaba2 P<sub>TEF1</sub>.bcaba3 P<sub>TEF1</sub>.bcaba4 P<sub>TEF1</sub>.bccpr1 P<sub>TEF1</sub>.tHMG1* | Verena Siewers, Chalmers University of Technology, Gothenburg, Sweden (3) |
|  | SABA3 | *S. cerevisiae* CEN.PK113-5D<br>*lpp1Δ::loxP dpp1Δ::loxP P<sub>ERG9Δ</sub>::loxP-P<sub>HXT1</sub> gdh1Δ::loxP P<sub>TEF1</sub>.ERG20 P<sub>PGK1</sub>.GDH2 P<sub>TEF1</sub>.tHMG1 P<sub>PGK1</sub>.bcaba1 P<sub>PGK1</sub>.bcaba2 P<sub>TEF1</sub>.bcaba3 P<sub>TEF1</sub>.bcaba4 P<sub>TEF1</sub>.bccpr1* | Verena Siewers, Chalmers University of Technology, Gothenburg, Sweden (3) |
|  | SbP-ABA1 | SbP<br>*P<sub>TDH3</sub>.bcaba1 P<sub>TEF1</sub>.bcaba2 P<sub>TDH3</sub>.bcaba3 P<sub>TEF1</sub>.bcaba4* | This study |
|  | E-ABA1 | Enterol<br>*P<sub>TDH3</sub>.bcaba1 P<sub>TEF1</sub>.bcaba2 P<sub>TDH3</sub>.bcaba3 P<sub>TEF1</sub>.bcaba4* | This study |
|  | E-ABA2 | Enterol<br>*P<sub>TDH3</sub>.bcaba1 P<sub>TEF1</sub>.bcaba2 P<sub>TDH3</sub>.bcaba3 P<sub>TEF1</sub>.bcaba4 P<sub>TDH3</sub>.bcaba1 P<sub>TDH3</sub>.bcaba3* | This study |
|  | E-ABA3 | Enterol<br>*P<sub>TDH3</sub>.bcaba1 P<sub>TEF1</sub>.bcaba2 P<sub>TDH3</sub>.bcaba3 P<sub>TEF1</sub>.bcaba4 P<sub>TEF1</sub>.bcaba1 P<sub>TEF1</sub>.bcaba3* | This study |
|  | E-ABA4 | Enterol<br>*P<sub>TDH3</sub>.bcaba1 P<sub>TEF1</sub>.bcaba2 P<sub>TDH3</sub>.bcaba3 P<sub>TEF1</sub>.bcaba4 P<sub>RPL18B</sub>.bcaba1 P<sub>RPL18B</sub>.bcaba3* | This study |
|  | E-ABA5 | Enterol<br>*P<sub>TDH3</sub>.bcaba1 P<sub>TEF1</sub>.bcaba2 P<sub>TDH3</sub>.bcaba3 P<sub>TEF1</sub>.bcaba4 P<sub>SAC6</sub>.bcaba1 P<sub>SAC6</sub>.bcaba3* | This study |
|  | E-CABA1 | Enterol<br>*P<sub>TDH3</sub>.bcaba1 P<sub>TEF1</sub>.bcaba2 P<sub>TDH3</sub>.bcaba3 P<sub>TEF1</sub>.bcaba4 P<sub>TEF1</sub>.bccpr1* | This study |
|  | E-CABA2 | Enterol<br>*P<sub>TDH3</sub>.bcaba1-bccpr1 P<sub>TEF1</sub>.bcaba2 P<sub>TDH3</sub>.bcaba3 P<sub>TEF1</sub>.bcaba4* | This study |
|  | E-CABA3 | Enterol<br>*P<sub>TDH3</sub>.bcaba1 P<sub>TEF1</sub>.bcaba2-bccpr1 P<sub>TDH3</sub>.bcaba3 P<sub>TEF1</sub>.bcaba4* | This study |
|  | E-CABA4 | Enterol<br>*P<sub>TDH3</sub>.bcaba1-bmr P<sub>TEF1</sub>.bcaba2 P<sub>TDH3</sub>.bcaba3 P<sub>TEF1</sub>.bcaba4* | This study |
|  | E-CABA5 | Enterol<br>*P<sub>TDH3</sub>.bcaba1 P<sub>TEF1</sub>.bcaba2-bmr P<sub>TDH3</sub>.bcaba3 P<sub>TEF1</sub>.bcaba4* | This study |
|  | E-CABA6 | Enterol<br>*P<sub>TDH3</sub>.bcaba1-bmr P<sub>TEF1</sub>.bcaba2-bccpr1 P<sub>TDH3</sub>.bcaba3 P<sub>TEF1</sub>.bcaba4* | This study |
|  | E-CABA7 | Enterol<br>*P<sub>TDH3</sub>.bcaba1-bccpr1 P<sub>TEF1</sub>.bcaba2-bmr P<sub>TDH3</sub>.bcaba3 P<sub>TEF1</sub>.bcaba4* | This study |
|  | E-TABA2 | Enterol<br>*P<sub>TDH3</sub>.bcaba1 P<sub>TEF1</sub>.bcaba2 P<sub>TDH3</sub>.bcaba3 P<sub>TEF1</sub>.bcaba4 P<sub>TDH3</sub>.bcaba1 P<sub>TDH3</sub>.bcaba3 P<sub>TEF1</sub>.tHMG1* | This study |
|  | E-TABA3 | Enterol<br>*P<sub>TDH3</sub>.bcaba1 P<sub>TEF1</sub>.bcaba2 P<sub>TDH3</sub>.bcaba3 P<sub>TEF1</sub>.bcaba4 P<sub>TEF1</sub>.bcaba1 P<sub>TEF1</sub>.bcaba3 P<sub>TEF1</sub>.tHMG1* | This study |
|  | E-TABA4 | Enterol<br>*P<sub>TDH3</sub>.bcaba1 P<sub>TEF1</sub>.bcaba2 P<sub>TDH3</sub>.bcaba3 P<sub>TEF1</sub>.bcaba4 P<sub>RPL18B</sub>.bcaba1 P<sub>RPL18B</sub>.bcaba3 P<sub>TEF1</sub>.tHMG1* | This study |
| | E-TABA5 | Enterol<br>$P_{TDH3}.bcaba1$ $P_{TEF1}.bcaba2$ $P_{TDH3}.bcaba3$ $P_{TEF1}.bcaba4$<br>$P_{SAC6}.bcaba1$ $P_{SAC6}.bcaba3$ $P_{TEF1}.tHMG1$ | This study |
| | E-TCABA2 | Enterol<br>$P_{TDH3}.bcaba1$ $P_{TEF1}.bcaba2$ $P_{TDH3}.bcaba3$ $P_{TEF1}.bcaba4$<br>$P_{TDH3}.bcaba1$ $P_{TDH3}.bcaba3$ $P_{TEF1}.tHMG1$ $P_{TEF1}.bccpr1$ | This study |
| | E-TCABA3 | Enterol<br>$P_{TDH3}.bcaba1$ $P_{TEF1}.bcaba2$ $P_{TDH3}.bcaba3$ $P_{TEF1}.bcaba4$<br>$P_{TEF1}.bcaba1$ $P_{TEF1}.bcaba3$ $P_{TEF1}.tHMG1$ $P_{TEF1}.bccpr1$ | This study |

Subsequently, we compared ABA production levels in the E-ABA1 strain to those in one of the best ABA-producing *S. cerevisiae* strains generated by Otto *et al.* (TABA3) (50). Our results show that after 12 h of cultivation at 30°C, the E-ABA1 strain produced significantly less ABA than the TABA3 strain. However, by 24 h, there was no significant difference in ABA production between the two strains (**Figure 1B**). While the TABA3 strain reached its maximum ABA production levels within 12 h, the E-ABA1 strain required 24 h to achieve similar production levels. This suggests that, despite having fewer integrated pathway genes, the E-ABA1 strain was able to produce ABA levels comparable to the TABA3 strain after 24 h.

### Expression of heterologous cytochrome P450 reductases further increases ABA production titers

A well-established strategy to enhance ABA production yield involves optimizing the expression of cytochrome P450 monooxygenases (CYPs) (55,56). In *S. cerevisiae* strains engineered to produce ABA via a heterologous pathway, the CYPs BcABA1 and BcABA2 have been identified as critical bottlenecks. Overexpression of CYPs can increase the demand for NADP^+^ and thereby potentially causes metabolic imbalances within the organism. These imbalances can be mitigated by co-overexpression of cytochrome P450 reductases (CPRs), which provide the necessary NADP^+^ equivalents (56). Otto *et al.* demonstrated that the CPR from *B. cinerea* (*bccpr1*) was more effective in enhancing ABA production in *S. cerevisiae* than the native *cpr* gene (50). Building on this finding, we evaluated the effect of integrating a copy of *bccpr1* into the E-ABA1 genome (**Table 1**). We constructed the E-CABA1 strain and showed that after 24 h of cultivation at 30°C in 3 mL of YPD, the inclusion of *bccpr1* resulted in a 7-fold increase in ABA titers in the supernatant (1.3 mg/L in E-ABA1; 9.7 mg/L in E-CABA1) (**Figure 2A**).

**Figure 2.**
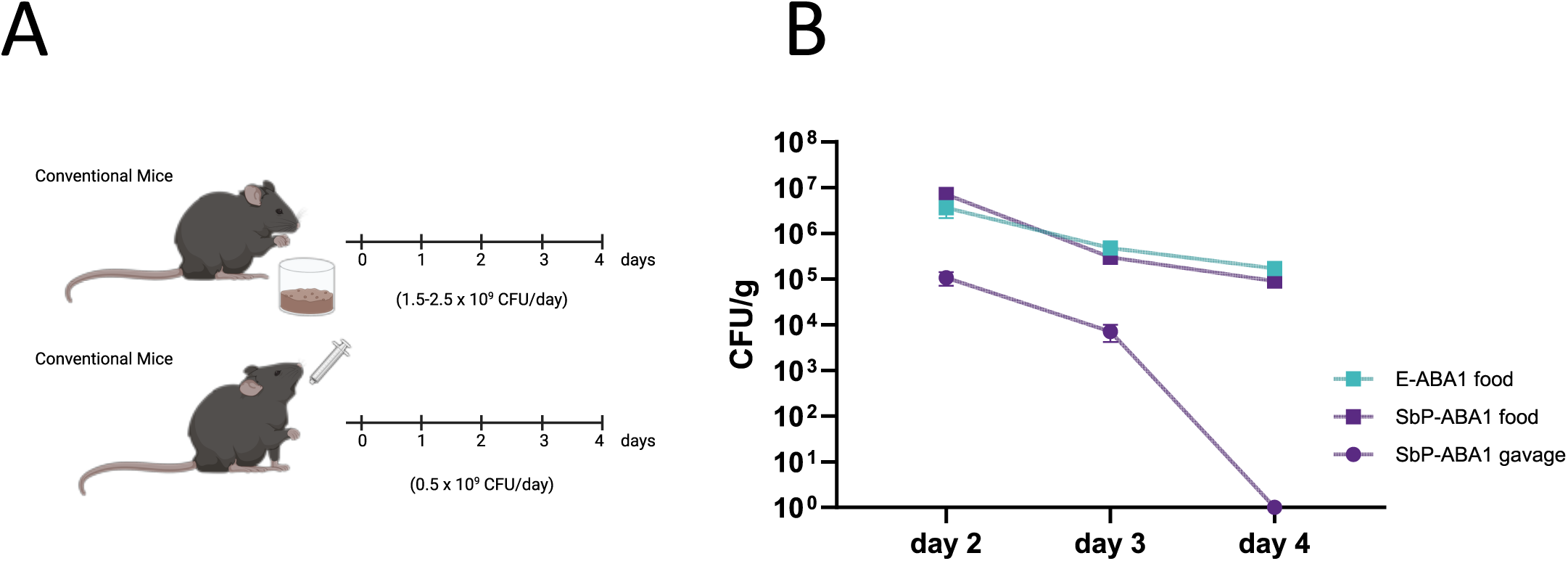
ABA production in E-ABA strains containing a heterologous cytochrome P450 reductase gene. **(A-B)** All strains were based on the E-ABA1 strain, which in an Enterol strain containing *bcaba1-4*. Average ABA titers were calculated from 4-6 independent biological replicates. Effect of *bccpr1* (E-CABA1), fusion of *bccpr1* to *bcaba1* (E-CABA2) or *bcaba2* (E-CABA3), fusion of *bmrBM3* to *bcaba1* (E-CABA4) or *bcaba2* (E-CABA5) and combined fusion of *bccpr1* to *bcaba1* and *bmrBM3* to *bcaba2* (E-CABA6) or *bccpr1* to *bcaba2* and *bmrBM3* to *bcaba1* (E-CABA7) on ABA titers. **(A)** Strains were cultivated for 24 h at 30°C in 3 mL of YPD medium in 24-well deep well plates. **(B)** Strains were cultivated for 24 h at 30°C in 3 mL of YPD medium in test tubes. Data are represented as mean ± SEM and statistically analyzed using a one-way ANOVA. * p < 0.05, ** p < 0.01, *** p < 0.001, **** p < 0.0001.

Furthermore, multiple reports have indicated that engineering redox partner fusions can support CYP activity by reducing complexity of the P450 system and improving electron transfer properties, thereby enhancing catalytic performance and redirecting flux towards particular pathway branches (9,57). We hypothesized that fusing *bccpr1* to the CYPs *bcaba1* and *bcaba2* could further increase ABA production. The corresponding strains were named E-CABA2 and E-CABA3, respectively (**Table 1**). Surprisingly, no significant differences in ABA titers were observed between E-ABA1 and E-CABA2 or E-CABA3 (**Figure 2A**). In addition to the *bccpr1* fusion, we evaluated an alternative reductase, BMR, from *Bacillus megaterium* BM3. This reductase is naturally fused to CYP enzymes and its fusion has been shown to dramatically increase CYP activity in other metabolic engineering applications (58,59). Therefore, we hypothesized that BMR could be better suited as a fusion partner. The strains where *bmrBM3* was fused to either *bcaba1* or *bcaba2* were named E-CABA4 and E-CABA5, respectively (**Table 1**). Consistent with the results of the bc*cpr1* fusion, fusing *bmrBM3* to either *bcaba1* or *bcaba2* did not result in significant changes in ABA expression levels (**Figure 2B**). Additionally, we also explored whether the titer could be increased by fusing both *cyp*s to a *cpr* gene. To this end, *bcaba1* was fused to *bmrBM3* and *bcaba2* to *bccpr1*, and vice versa, resulting in the strains E-CABA6 and E-CABA7, respectively (**Table 1**). No big differences in OD_600_ were observed between E-ABA1 and the strains containing additional *cpr*s (**Figure S1B,C**). After 24 h of cultivation at 30°C in 3 mL of YPD medium, we observed a 2.5-fold increase in ABA production in the strains with a CPR fused to both *bcaba1* and *bcaba2* compared to E-ABA1 (E-ABA1 = 1.3 mg/L; E-CABA6 = 3.2 mg/L; E-CABA7 = 3.4 mg/L) (**Figure 2A**). Nonetheless, the increase in ABA production observed for CPR-fusions was still lower than the 7-fold increase observed when *bccpr1* was integrated separately.

### Analysis of mouse gut yeast colonization and mouse serum ABA levels

To ensure effective ABA delivery to the gut, the engineered yeast strain must be able to produce ABA not only in culture but also within the gut environment. Considering the significant differences between *in vitro* culture conditions and the gut, we evaluated the strain’s ability to colonize the gut and measured serum ABA levels in mice.

To administer yeast strains to mice, oral gavage is a commonly used method. However, since daily oral gavage is known to induce considerable stress in animals, we developed an optimized, less stressful approach for yeast delivery. In this method, yeast was incorporated into the mice’s diet. The yeast-supplemented diet was prepared by mixing standard powdered mouse food with water and yeast pellets to form a porridge-like consistency (**Figure S2A**).This mixture was placed into glass jars secured in stainless steel holders to prevent moving or tipping over of the jars (**Figure S2B**). The feeding system was then placed into the cages, allowing *ad libitum* consumption (**Figure S2C**). In preliminary tests comparing various yeast concentrations per gram of food, we observed that mice showed no preference for low or high concentrations. As a result, we conducted colonization experiments using the highest concentration, with 5 x 10^8^ colony forming units (CFU) added per gram of food. Since mice typically consume 3-5 grams per day, this corresponded to a daily dose of approximately 1.5-2.5 x 10^9^ CFU. In the oral gavage set-up, a lower dose of yeast (5 x 10^8^ CFU) was administered due to limitations in the amount of yeast pellet that could be dissolved within a volume suitable for gavage.

To assess the ability of yeast strains to colonize the gut, mice were administered an engineered yeast for three consecutive days, either through a yeast-supplemented diet or through oral gavage with strains E-ABA1 or SbP-ABA1 (experimental design in **Figure 3A**). Yeast presence was monitored in the stool via CFU counting. While high levels of yeast were still detectable in the stool two days after the last feeding with yeast-supplemented diet, yeast was undetectable in the stool of gavaged mice after the same period (**Figure 3B**).

**Figure 3.**
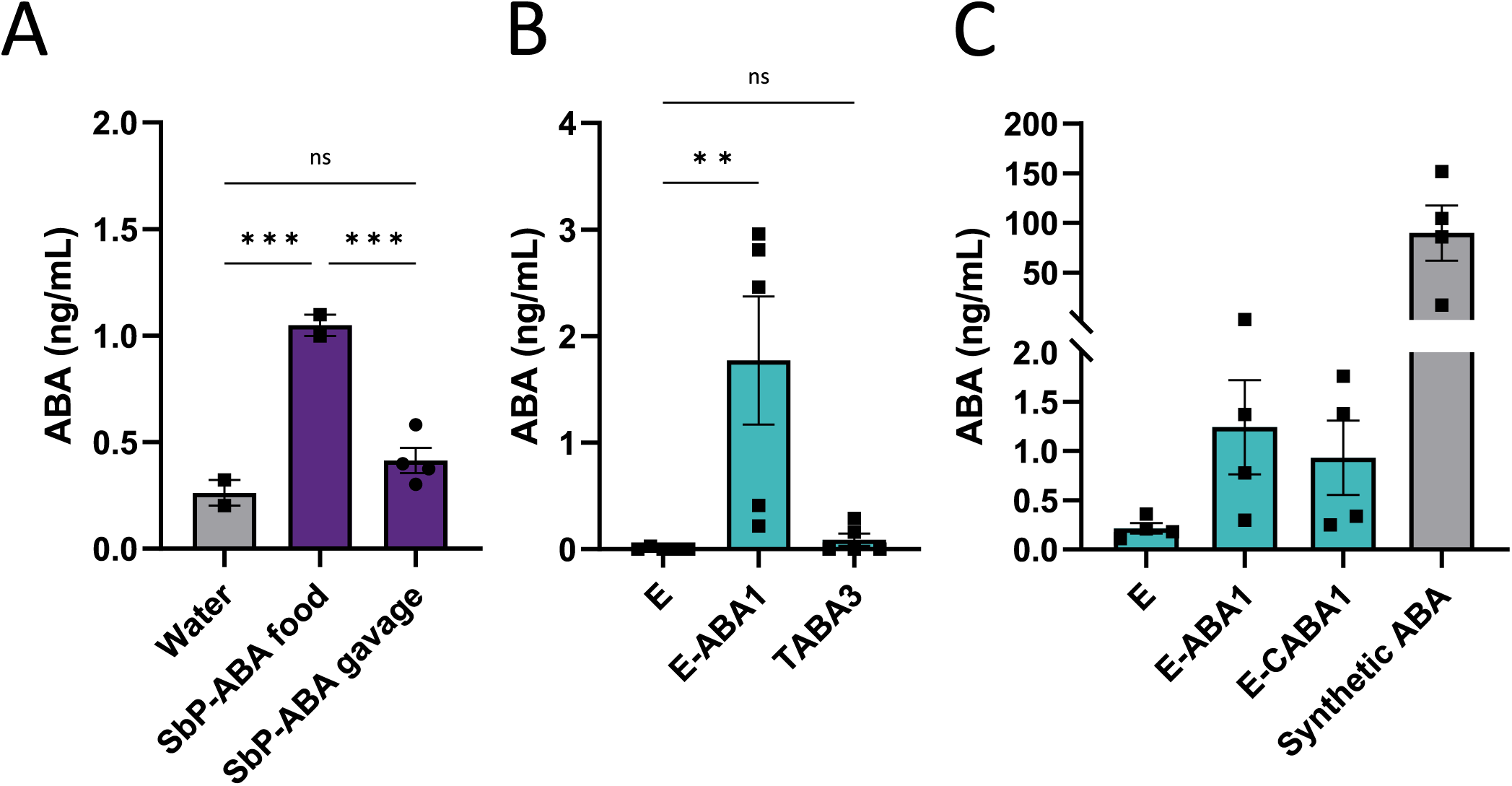
Determination of CFU count in the feces of conventional mice that received either the yeast supplemented diet or were administered the yeast through oral gavage. **(A)** Schematic representation showing treatment of the mice. First, conventional mice were fed a yeast-supplemented diet containing 5 x 10^8^ CFU/g of food. The diet included either E-ABA1 or SbP-ABA1 and was administered for 3 consecutive days (d 0, d 1, d 2) (yellow dots). Second, conventional mice received 3 doses of 5 x 10^8^ CFU of SbP-ABA1 through oral gavage (d 0, d 1, d 2) (yellow dots). The time points for collection of feces are depicted by brown dots. **(B)** Fecal carriage of yeast in both set-ups. Fecal samples were collected as shown in the timelines, plated on YPD media containing chloramphenicol (50 mg/L) and kept at 30°C for 2 d. Data are represented as mean ± SEM (n = 3 mice). Abbreviations: E-ABA1 = Enterol strain containing *bcaba1-*4, SbP-ABA1 = SbP strain containing *bcaba1-*4, CFU = colony forming units.

Additionally, we measured ABA levels in the serum of mice following 1 week of treatment with either a yeast-supplemented diet or yeast administered via oral gavage. While oral gavage of the SbP-ABA1 strain did not result in an increase in serum ABA levels (**Figure 4A**), mice fed the SbP-ABA1 and E-ABA1 strains exhibited serum ABA levels of 1.0 ng/mL and 1.8 ng/mL, respectively, both of which were significantly higher than the levels observed in mice fed the wild-type Enterol strain or a regular diet (**Figure 4A,B**). Interestingly, this increase in serum ABA was not observed in mice treated with the most productive ABA-producing *S. cerevisiae* strain, TABA3 (50) (**Figure 4B**). However, it is important to note that the CFU dose administered to these mice was 2-fold lower due to slower growth rate of this strain. Despite the higher ABA production observed in YPD cultures for the E-CABA1 strain compared to E-ABA1 (**Figure 2A**), no corresponding increase in serum ABA levels was detected in mice receiving this strain (**Figure 4C**). Furthermore, the ABA levels in mice fed E-ABA1 were approximately 70 times lower than those observed in mice treated with 0.3-0.5 mg of ABA via a diet supplemented with 100 mg/kg of synthetic ABA (**Figure 4C**).

**Figure 4.**
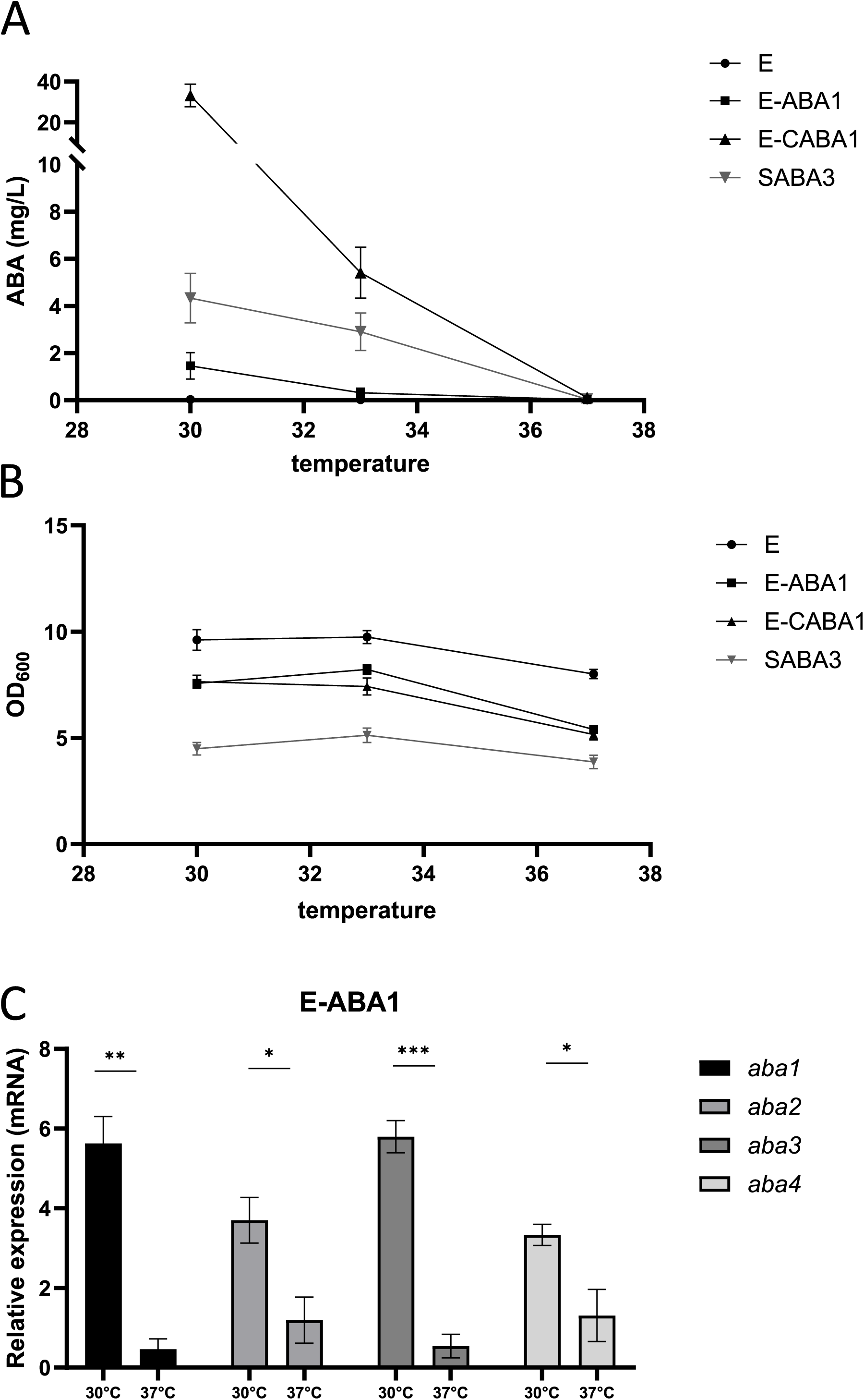
ABA titers in serum of mice fed or gavaged with an ABA-producing yeast for 1 week. **(A)** Mice were either fed a yeast-supplemented diet containing ABA-producing strain SbP-ABA1 (5 x 10^8^ CFU/g of food) or received oral gavage with this strain (5 x 10^8^ CFU). SbP-ABA1 = SbP strain containing *bcaba1-4*. **(B)** Mice were fed a yeast-supplemented diet containing either wild-type Enterol (E) (5 x 10^8^ CFU/g of food), the ABA-producing Enterol strain E-ABA1 (containing *bcaba1-4*) (5 x 108 CFU/g of food) or the ABA-producing *S. cerevisiae* strain TABA3 (containing *bcaba1-4*, *bccpr1*, *tHMG1*) (2 x 10^8^ CFU/g of food). **(C)** Mice were fed a yeast-supplemented diet containing either wild-type Enterol (5 x 10^8^ CFU/g of food), the ABA-producing Enterol strains E-ABA1 (5 x 10^8^ CFU/g of food) or E-CABA1 (containing *bcaba1-4*, *bccpr1*) (5 x 10^8^ CFU/g of food) or a diet supplemented with synthetic ABA (100 mg/kg of food). Serum was taken at sacrifice after 1 week of treatment. Each dot represents one mouse. Data are represented as mean ± SEM and statistically analyzed using a one-way ANOVA. * p < 0.05, ** p < 0.01, *** p < 0.001, **** p < 0.0001.

Given that ABA was detected in the serum of mice fed the yeast-supplemented diet but not in mice treated via oral gavage, it remained unclear whether the detected ABA was produced by the yeast when in the gut or synthesized by the yeast in the feedings bowl prior to ingestion. To investigate this, we kept feeding bowls containing yeast-supplemented food at room temperature for 0, 8 or 24 h. To measure ABA levels, water was added to the food to create a diluted, homogeneous suspension, and ABA was then quantified in the supernatant. We found that ABA was detectable in feeding bowls stored at room temperature for 24 h (**Figure S3A**). This duration corresponds to the time the feeding bowls remained in the cage before being refreshed during the mouse experiment, suggesting that at least some of the ABA detected in serum may have been produced in the feeding bowls rather than within the gut. We hypothesized that a low or absent ABA production within the gut could be a result of genetic variations arising during gastrointestinal transit or competitive interactions with the microbiome. To investigate whether the absence of ABA production was due to genetic alterations in the yeast after passing through the gastrointestinal tract, stool pellets from mice fed the yeast-supplemented diet were plated on agar plates and individual yeast colonies were isolated and cultured in liquid media at 30°C for 24 h. ABA was detected in the supernatant of these cultures, indicating that the yeast retained its ability to produce ABA after passage through the gut (**Figure S3B**). To explore whether competition with the native gut microbiota affected ABA production, germ-free mice were gavaged with the engineered ABA-producing yeast strain E-ABA1 (experimental design in **Figure S4A**). The mice received a single dose of E-ABA1 on d 0 and the presence of this yeast was assessed in stool samples on d 3, 7 and 13 post-gavage. ABA levels were measured in serum on d 13 post-gavage. Despite detecting high concentrations of E-ABA1 in the stool (**Figure S4B**), ABA remained undetectable in the serum (data not shown), indicating that microbial competition was not the cause for low ABA levels in serum.

### Identification of physiological body temperature (37°C) as ABA production bottleneck

The engineered yeast successfully colonized the gut (**Figure 3B**) and remained both viable and genetically stable following passage through the gastrointestinal tract (**Figure S3B**). However, we were unable to detect ABA in serum of mice that received an oral gavage with this yeast (**Figure 4A**). To investigate this limitation, the yeast strain’s ability to produce ABA at various culturing temperatures was evaluated. Three specific temperatures were selected for this assessment: 30°C, 33°C and 37°C, representing *in vitro* growth conditions, an intermediate measurement, and the physiological temperature of mice, respectively. ABA production was compared between E-ABA1, E-CABA1 and the SABA3 *S. cerevisiae* strain of Otto *et al.* (50) (**Figure 5A**). The SABA3 strain, in addition to the modifications made in the previously described TABA3 strain, was intensively modified to enhance upstream flux through the MVA pathway and to reduce the formation of side products, thereby boosting the flux towards ABA synthesis. All yeasts in this experiment were pre-cultured at 30°C and subsequently adjusted to an OD_600_ of 0.2 before being incubated for 24 h at 30°C, 33°C or 37°C. At 30°C, the SABA3 strain demonstrated higher ABA production than the E-ABA1 strain, whereas the E-CABA1 strain exhibited seven times higher ABA production than the SABA3 strain. At 33°C, ABA production in the E-ABA1 strain was nearly abolished, and the E-CABA1 strain showed a 6-fold reduction in ABA production. In contrast, ABA levels in the SABA3 strain remained relatively stable under the same conditions, exhibiting only a 1.5-fold decrease. At 37°C, ABA production was completely abrogated in all strains tested. This reduction or loss of ABA production was not linked to decreased growth, as OD_600_ values at 33°C were comparable to those at 30°C. Although a reduction in OD_600_ was observed at 37°C, it was similar to that observed in the wild-type Enterol strain, suggesting that the reduced growth was not attributable to the introduced genetic modifications (**Figure 5B**).

**Figure 5.**
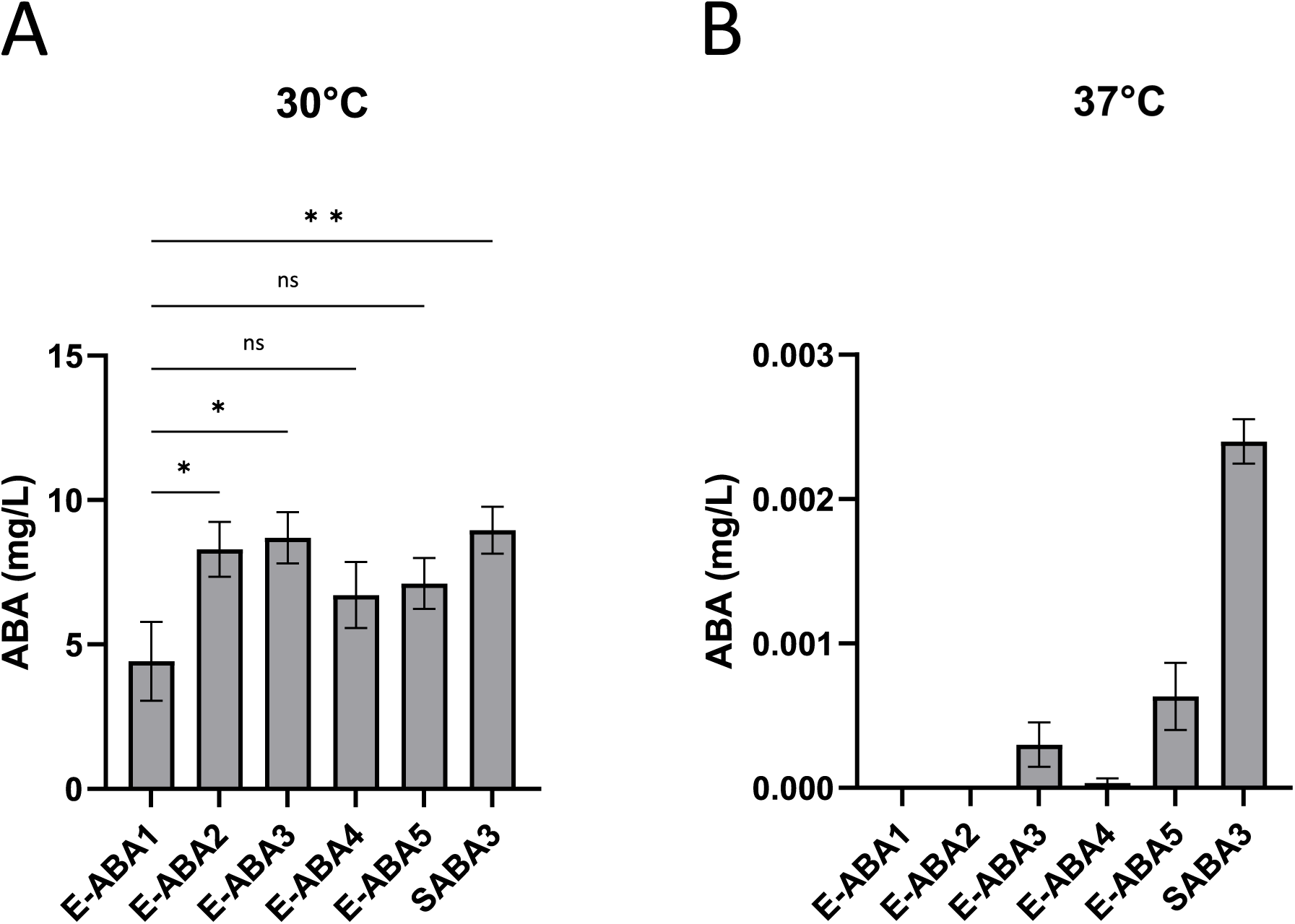
Effect of temperature on ABA production levels and mRNA expression levels of *bcaba1-4*. **(A)** ABA titer in supernatant of strains Enterol (E), E-ABA1, E-CABA1 and SABA3 (50). E-ABA1 and E-CABA1 are based on the genetic background of *S. boulardii* strain Enterol and contain *bcaba1-4*, and *bcaba1-4* and *bccpr1* respectively. SABA3 (50) is based on *S. cerevisiae* strain CENK.PK113-5D and contains various modifications as shown in **Table 1**. Strains were cultivated for 24 h in 3 mL of YPD medium in 24-well deep well plates. Average ABA titers were calculated from 3 independent biological replicates and error bars indicate the standard error of the mean. **(B)** OD_600_ of strains analyzed in Figure 5A. **(C)** mRNA expression of *bcaba1*, *bcaba2*, *bcaba3* and *bcaba4* normalized to expression of *25S rRNA*, measured in the pellet of a 24 h culture of E-ABA1 grown at 30°C or 37°C in 3 mL of YPD in test tubes. Average expression was calculated from 3 independent biological replicates. Data are represented as mean ± SEM and statistically analyzed using a Student’s t-test. * p < 0.05, ** p < 0.01, *** p < 0.001, **** p < 0.0001.

### Effect of promoter strength on ABA production at physiological body temperature

Given the decrease in ABA production at higher temperatures, qPCR was conducted to compare the expression levels of the different *aba* genes in the heterologous pathway at both low (30°C) and high (37°C) temperatures. This analysis aimed to identify potential bottleneck genes in the pathway. We observed that the expression of all four *aba* genes was significantly reduced at 37°C compared to at 30°C (**Figure 5C**). Specifically, the expression levels of *bcaba2* and *bcaba4* decreased by 3-fold and 2.5-fold, respectively, while *bcaba1* and *bcaba3* exhibited a more pronounced decrease of 12-fold and 11-fold, respectively. Notably, *bcaba1* and *bcaba3* were integrated into the genome in a bicistronic manner under the *TDH3* promoter, while *bcaba2* and *bcaba4* were expressed in a bicistronic manner under the *TEF1* promoter. A similar effect was observed in the TABA3 strain, where genes regulated by the *TEF1* promoter (*bcaba3* and *bcaba4*) exhibited a 2-fold decrease in expression levels, while those controlled by the *PGK1* promoter (*bcaba1* and *bcaba2*) showed a more pronounced, 4-fold reduction in expression (**Figure S5**). This prompted us to hypothesize that the reduction in ABA production at 37°C might be a result of decreased promoter activity at higher temperatures. To investigate this, we introduced an additional copy of the *bcaba1-P2A-bcaba3* expression cassette under various promoters with different expression strengths, including *TDH3*, *TEF1*, *RPL18B* and *SAC6* (35), resulting in the creation of strains E-ABA2, E-ABA3, E-ABA4 and E-ABA5, respectively (**Table 1**). At 30°C, the inclusion of an extra copy of *bcaba1-P2A-bcaba3* under the control of the *TDH3*, *TEF1* or *SAC6* promoter led to increased expression of these genes, whereas the *RPL18B* promoter did not show the same effect (**Figure S6A,B**). At 37°C, an increase in *bcaba1* and *bcaba3* gene expression was observed with the additional copy under the control of the *TEF1*, *RPL18B* or *SAC6* promoters, but not under the *TDH3* promoter (**Figure S6A,B**). Introduction of an additional copy of *bcaba1-P2A-bcaba3* did not result in changes is OD_600_ (**Figure S6C**). The integration of an additional copy of these genes significantly increased ABA production titers at 30°C for the strong promoters *TDH3* and *TEF1*, with a trend towards increased ABA titers for the intermediate and weak promoters *RPL18B* and *SAC6* (**Figure 6A**). However, at 37°C ABA titers remained near zero for all strains, regardless of the promoter used (**Figure 6B**). Therefore, decreased promoter activity was not identified as the primary cause of the abrogated ABA production at 37°C.

**Figure 6.**
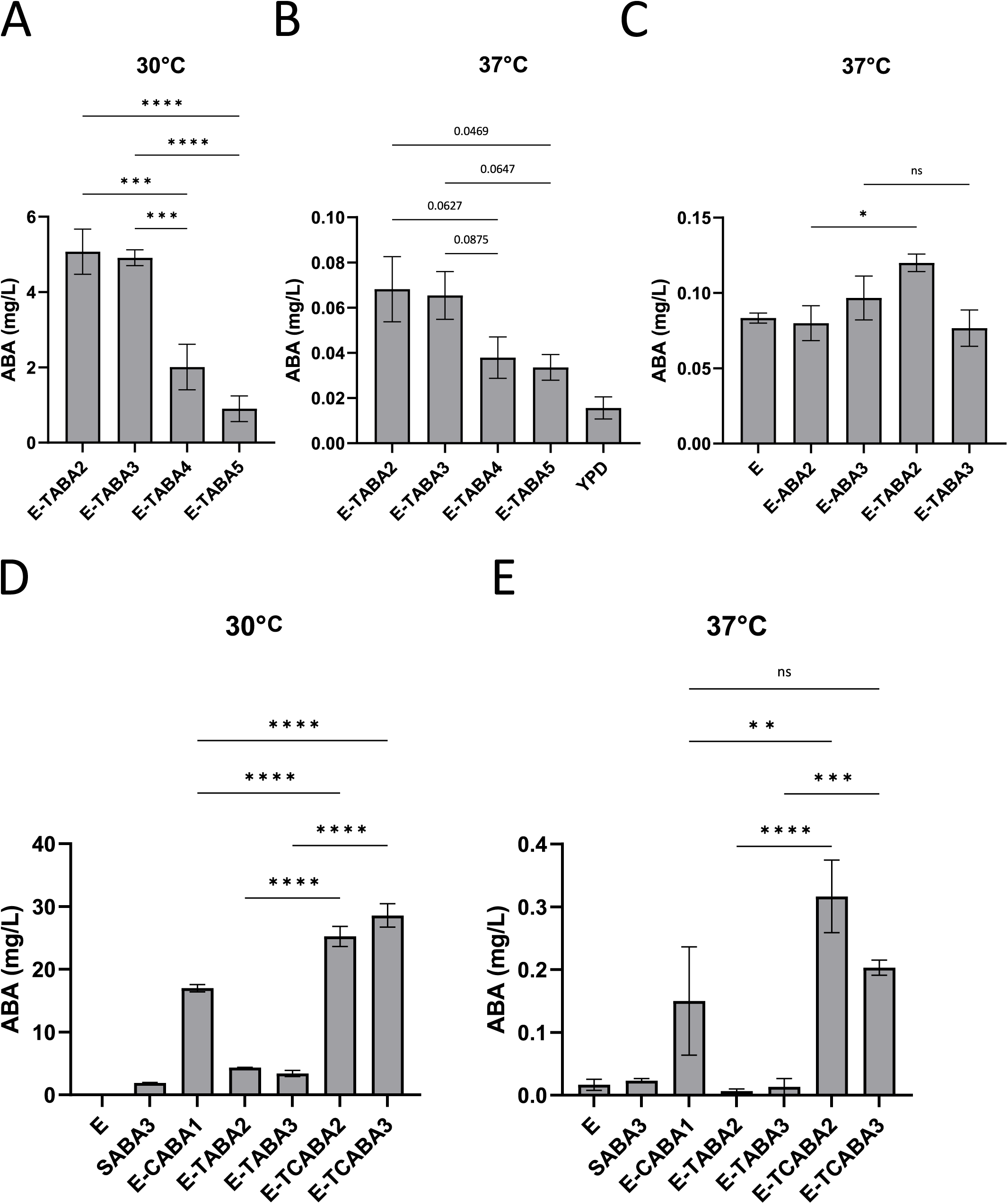
ABA production in E-ABA strains containing an additional copy of *bcaba1* and *bcaba3* genes expressed under various promoters. All described strains were based on the E-ABA1 strain (Enterol containing *bcaba1-4*). Average ABA titers were calculated from 3 independent biological replicates. Effect of an additional copy of *bcaba1-P2A-bcaba3* expressed under control of the *TDH3* (E-ABA2), *TEF1* (E-ABA3), *RPL18B* (E-ABA4) or *SAC6* (E-ABA5) promoter on ABA titers. SABA3 (50), based on *S. cerevisiae* strain CENK.PK113-5D and contains various modifications as shown in **Table 1**, is included as a control. Strains were cultivated for 24 h in 3 mL of YPD medium in 24-well deep well plates. **(A)** Cultures were grown at 30°C. **(B)** Cultures were grown at 37°C. Data are represented as mean ± SEM and statistically analyzed using a one-way ANOVA. * p < 0.05, ** p < 0.01, *** p < 0.001, **** p < 0.0001.

### Generation of a yeast strain producing low levels of ABA at 37°C

Recognizing the need to achieve high production titers for the therapeutic application of an ABA-producing probiotic, we explored various strategies to further enhance ABA yields. Given the stability of ABA production at 33°C compared to 30°C in the *S. cerevisiae* strain SABA3, which is heavily boosted in MVA flux, we hypothesized that increasing MVA flux in our Enterol strains could similarly boost ABA production at higher temperatures. To test this hypothesis, we integrated truncated 3-hydroxy-3-methylglutaryl-CoA reductase (*tHMG1)*, coding for a rate-limiting enzyme in the MVA pathway, into the genome. This modification was introduced into the strains with an additional copy of *bcaba-P2A-bcaba3,* resulting in the creation of strains E-TABA2, E-TABA3, E-TABA4 and E-TABA5 (**Table 1**). Our results showed that at 30°C, E-TABA2 and E-TABA3 exhibited 2.5-fold and 5.5-fold higher ABA production compared to E-TABA4 and E-TABA5, respectively (**Figure 7A**). Although ABA levels were lower at 37°C, E-TABA2 and E-TABA3 still produced twice as much ABA as E-TABA4 and E-TABA5 (**Figure 7B**). E-TABA2 and E-TABA3 were identified as the highest-producing strains containing *tHMG1,* and E-TABA2 in particular demonstrated a statistically significant, 1.5-fold, increase in ABA titers compared to E-ABA2 (**Figure 7C**). To further enhance ABA yield, we introduced a copy of *bccpr1,* which encodes a cytochrome P450 reductase and is known from previous experiments to substantially increase ABA titers (**Figure 2A**). The resulting strains, named E-TCABA2 and E-TCABA3 (**Table 1**), showed the highest ABA production levels at 30°C among all strains tested (25.2 mg/L in E-TCABA2; 28.6 mg/L in E-TACABA3), significantly outperforming SABA3, E-CABA1 and their E-TABA counterparts (1.9 mg/L in SABA3; 17 mg/L in E-CABA1; 4.3 mg/L in E-TABA2, 3.4 mg/L in E-TABA3) (**Figure 7D**). At 37°C, these strains produced less than 0.5 mg/L of ABA, but for the first time, ABA levels showed a more than 10-fold increase compared to background ABA levels measured in wild-type Enterol (0.32 mg/L in E-TCABA2; 0.20 mg/L in E-TCABA3; 0.02 mg/L in E) (**Figure 7E**). Notably, E-TCABA2 and E-TCABA3 also produced significantly higher ABA titers than their E-TABA counterparts (0 mg/L in E-TABA2; 0.01 mg/L in E-TABA3), with E-TCABA2 outperforming E-CABA1 (0.15 mg/L in E-CABA1), as was also observed at 30°C. Consequently, we successfully generated a strain capable of producing low amounts of ABA at 37°C, with levels equal to or higher than 0.2 mg/L. This represents a significant increase compared to production levels measured in previously generated strains.

**Figure 7.**
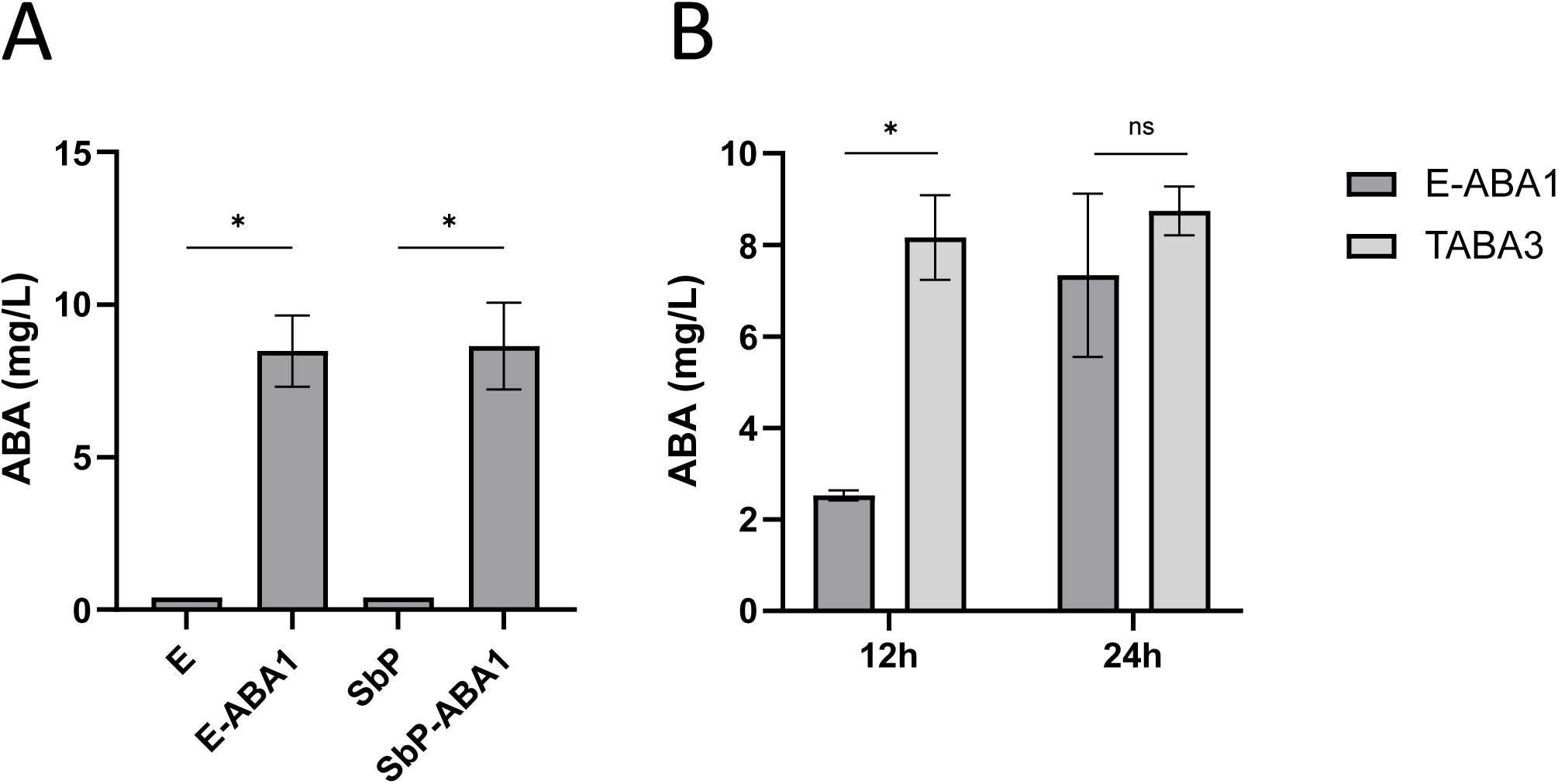
ABA production in E-ABA strains containing a copy of *tHMG1* and/or *bccpr1*. **(A-C)** The effect of adding *tHMG1* to E-ABA2, E-ABA3, E-ABA4 and E-ABA5 on ABA titers during cultivation at **(A)** 30°C or **(B, C)** 37°C. More information on genetic composition of strains can be found in **Table 1**. **(D-E)** The effect of adding *bccpr1* to E-TABA2 and E-TABA3 on ABA titers during cultivation at **(D)** 30°C or **(E)** 37°C. More information on genetic composition of strains can be found in **Table 1**. E = wild-type Enterol; YPD = blank YPD medium. SABA3 = S. cerevisiae strain contains various modifications (50) as shown in **Table 1**. Strains were cultivated for 24 h in 3 mL of YPD medium in 24-well deep well plates. Average ABA titers were calculated from 3-5 independent biological replicates. Data are represented as mean ± SEM and statistically analyzed using a one-way ANOVA. * p < 0.05, ** p < 0.01, *** p < 0.001, **** p < 0.0001.

## Discussion

ABA is considered an interesting nutraceutical substance that may explain some of the health benefits associated with consumption of ABA-rich foods (2). Since challenges such as scalability, short half-life and cost-effectiveness hinder the practical use of ABA-enriched fruit extracts and synthetic ABA supplements (9,10), we propose an alternative ABA administration technique in this study by engineering an ABA-producing probiotic strain of *Saccharomyces boulardii*. Consistent with observations made in *S. cerevisiae* (50)*, Yarrowia lipolytica* (60) and *Aspergillus oryzae* (46), we demonstrated that the integration of the *bcaba1-4* expression cassette from *Botrytis cinerea* is sufficient to establish ABA production in *S. boulardii* (**Figure 1A**). Additionally, the subsequent incorporation of the *bccpr1* gene led to a 7-fold increase in ABA production (**Figure 2A**), and already surpassed the performance of the highest-producing *S. cerevisiae* strain previously generated by Otto *et al.* (50) (**Figure 5A**). To further increase ABA production, we engineered redox partner fusions, which are generally recognized for enhancing CYP activity by improving electron transfer efficiency and catalytic performance (9,56,57). In our study, fusing *bc*c*pr1* and *bmrBM3* to the CYP enzymes *bcaba1* and *bcaba2* resulted in only a 2.5-fold increase in ABA production, which was lower than the 7-fold increase observed when *bccpr1* was integrated separately (**Figure 2A**). Previous studies have indicated that the length of the linker is a critical determinant of enzyme activity (61) and that the activity of fusion constructs can be fine-tuned through linker engineering (57). Therefore, optimizing linker length remains a promising strategy to further enhance ABA production.

To develop a genetically engineered probiotic strain effectively delivering ABA to the gut, the strain should possess the following attributes: tolerance to low pH to ensure survival and genetic stability during gastrointestinal transit, the ability to adhere to the intestinal epithelium for effective colonization, and efficient ABA production at the epithelial cell surface (62).

We demonstrated that our ABA-producing *S. boulardii* remained both viable and genetically stable following passage through the gastrointestinal tract as colonies recovered from stool samples were able to produce ABA in YPD cultures at 30°C with titers comparable to those observed in the original cultures prior to ingestion (**Figure S3B**).

By feeding mice a yeast-supplemented diet, we showed that our ABA-producing *S. boulardii* effectively colonized the gut, as evidenced by a high number of CFU detected in the stool two days after the cessation of the yeast-supplemented diet (**Figure 3A,B**). In contrast, when mice received an oral gavage with a yeast dose that was three times lower than the amount received through the yeast-supplemented diet, the yeast was cleared from the gut more rapidly, with no detectable CFU two days after the last gavage. These results suggest that enhancing the gut adhesion capacity of *S. boulardii* could be beneficial. A potential strategy to improve gut adhesion involves genetic engineering to introduce or overexpress specific adhesion-related proteins. One approach could be the integration of the adhesin gene *epa1* from *Candida glabrata*, or the creation of a chimeric protein by combining the endogenous *S. boulardii flo1* homolog with the N-terminal carbohydrate-binding domain of *Cgepa1* (63). Additionally, other EPA proteins with different glycan-binding specificities could be explored to enhance gut colonization (64). Although *C. glabrata* is closely related to *Saccharomyces* (as *Candida* is not monophyletic), extensive genetic modifications would be required to confer pathogenic traits to *S. boulardii* (65), suggesting that this modification could be safe. However, introducing a pathogenicity-associated factor like EPA1 into a probiotic yeast such as *S. boulardii* still poses a potential risk that requires careful evaluation before clinical application. If proven successful and safe, this approach could potentially benefit a wide range of engineered probiotics based on *S. boulardii*.

Although successful gut colonization was achieved in mice fed the yeast-supplemented diet, we observed only minimal levels of ABA in their serum (**Figure 4A-C**). Notably, no ABA was detected when the yeast was administered via oral gavage (**Figure 4A**). Our findings suggest that the low serum ABA levels in mice fed the yeast-supplemented diet were due to ABA being produced in the feedings bowl at room temperature prior to ingestion, rather than within the gut at 37°C (**Figure S3A**). *In vitro* experiments confirmed that the physiological body temperature of 37°C is likely a critical factor limiting ABA production by the engineered *S. boulardii* within the gut (**Figure 5A**).

Recognizing the importance of achieving high ABA production titers for the therapeutic application of an ABA-producing probiotic, we explored several strategies to further enhance ABA yields at 37°C. Firstly, we considered the possible inactivation of the *Botrytis* ABA biosynthesis enzymes at temperatures above 30°C. However, our *in silico* predictions for the thermal stability of BcABA3 and BcABA4 did not suggest that these enzymes are particularly temperature-sensitive (66,67). However, due to their large size, BcABA1 and BcABA2 could not be analyzed with the *in silico* tool. Despite this, 3D modeling of these proteins indicates a complex network of interactions, suggesting no significant issues with their thermal stability.

Nevertheless, it could be worthwhile to explore alternative enzymes in the ABA biosynthesis pathway from other phytopathogenic fungi, such as *Leptosphaeria maculans* (68) and *Magnaporthe oryzae* (69). Secondly, the availability of molecular oxygen may also be a potential rate-limiting factor for ABA production in the gut, as two catalytic steps in the ABA biosynthesis pathway require oxidation. While the gut lumen is mostly anaerobic, there is an oxygen gradient, with oxygen levels near the gut wall being similar to those in the bloodstream (70). Therefore, yeasts that adhere efficiently to the gut epithelium might still access sufficient oxygen for ABA production. Decreased thermotolerance of the cells is also an important factor to consider in bioengineering (71). Thermotolerance, a polygenic trait, is crucial for cell survival and growth at high temperatures. Previous studies in *S. cerevisiae* have shown that increasing the gene copy number of heterologous CYPs and CPRs results in decreased growth rates, increased formation of reactive oxygen species (ROS) (72) and reduced resistance to other cellular stresses such as elevated temperature (60). This suggests that reduced thermotolerance might be contributing to the poor ABA production observed in our strain at 37°C. However, our observations indicated that cell survival at 37°C was not compromised, as measured by acridine orange counting (data not shown). Additionally, although the OD_600_ of ABA-producing *S. boulardii* was significantly lower at 37°C compared to 30°C after 24 h, this reduction was similar to the decrease observed in wild-type *S boulardii* under the same conditions (**Figure 5B**). Nonetheless, improving thermotolerance might be an interesting strategy for increasing ABA production at 37°C. One method to enhance thermotolerance is experimental evolution, where yeast are gradually exposed to higher temperatures (71). However, this process requires many generations to achieve successful adaptation (73). Alternatively, thermotolerance can be enhanced more rapidly through whole-genome transformation (74) or through introduction of specific mutations in genes known to be involved in sustaining high thermotolerance. Mutations in the ergosterol biosynthesis pathway (e.g. in the genes *erg3*, *erg4* and *erg5*) have been shown to improve membrane fluidity, cell viability, cell integrity and overall thermotolerance (73,75,76). Furthermore, another promising method to further enhance ABA yield is balancing the heterologous ABA production pathway with the yeast’s native metabolism. To maximize ABA production at 37°C, it is crucial to achieve high expression levels of the *bcaba1-4* genes while maintaining robust expression of the native metabolism. A well-documented and effective strategy for balancing pathways in *S. cerevisiae* involves modifying promoter strength to coordinate gene expression (77,78). Consistent with previous studies showing a strong reduction in promoter activity under stress conditions, including elevated temperature (79), we observed that the expression TDH3, which is the promoter driving *bcaba1-P2A-bcaba3*, led to a decrease in mRNA expression of 12- and 11-fold respectively at 37°C (**Figure 5C**). This reduction may account for the absence of ABA production at 37°C. In an attempt to counteract this decreased expression, we integrated an additional copy of the *bcaba1-P2A-bcaba3* expression cassete under promoters with varying strength. Interestingly, while *TEF1*-, *RPL18B*- and *SAC6*-driven expression of *bcaba1-P2A-bcaba3* was higher at 37°C than at 30°C, ABA production remained below 0.001 mg/L at 37°C in these strains (**Figure 6B**).

Consequently, decreased promoter activity was not identified as the primary cause of the abrogated ABA production at 37°C. Moreover, balancing the heterologous pathway may also be achieved through redirection of redox cofactors and reduction of the metabolic burden on the cells (80). Redox constraints are known to limit the yield of many fermentation processes (81,82) and rerouting of NADPH-dependent synthetic pathways has been used to enhance heterologous production in *S. cerevisiae* (83,84). However, since Arnesen *et al.* showed that improving NADPH supply did not significantly increase ABA production in *Y. lipolytica*, this strategy might not be the most promising for increasing ABA titers in our *S. boulardii strain* (60). Lastly, the ABA levels in the SABA3 strain, which is engineered to enhance the upstream flux through the MVA pathway, remained relatively stable at 33°C compared to 30°C, exhibiting only a 1.5-fold decrease. In contrast, the E-CABA1 strain showed a more pronounced reduction, with a 6.1-fold decrease in ABA levels at 33°C (**Figure 5A**). These findings suggested that optimizing the biosynthetic pathway upstream of FPP and minimizing the formation of side products could be a promising strategy to further increase ABA production at 37°C. In our most modified Enterol strain, containing *bcaba1-4*, an additional copy of *bcaba1-P2A-bcaba3*, a copy of *bccpr1* and *tHMG1,* we observed an approximately 15-fold increase in ABA levels at both 30°C and 37°C compared to the highest-producing strain generated by Otto *et al.* (50). At 37°C, ABA concentrations reached or exceeded 0.2 mg/L (**Figure 7E**).

In summary, we report that despite successful colonization and survival in the gut, ABA production by our *S. boulardii* strain was hindered and the physiological body temperature (37°C) was identified as a significant bottleneck. Since serum ABA levels remained very low in mice fed a yeast-supplemented diet, further modifications are required before this probiotic can be tested in disease contexts in mice. Suggested modifications include optimizing the upstream biosynthesis pathway and enhancing thermotolerance through random or targeted mutations. Once the issue of *in vitro* production at 37°C is resolved, *in vivo* ABA serum levels can be assessed and possibly further increased through modifications that enhance gut adhesion.

## Materials and methods

### Microorganisms

NEB® *5-alpha Competent E. coli* cells (New England Biolabs) were used for plasmid amplification. The *Saccharomyces cerevisiae* strains used in this study are listed in detail in **Table 1**.

### Media and culture conditions

Bacterial cells were propagated in LB medium containing 10 g/L tryptone, 5 g/L yeast extract, 10 g/L NaCl and 100 mg/L ampicillin at 37°C. For agar plates, 15 g/L bacto agar was added.

Yeast cells were propagated at 30°C, 33°C or 37°C in YPD medium containing 10 g/L yeast extract, 20 g/L bacteriological peptone and 20 g/L glucose. Solid nutrient plates were made with 15 g/L bacto agar. Transformed strains were selected on agar plates containing appropriate antibiotics i.e. 200 mg/L geneticin (G418) or 100 mg/L nourseothricin.

### Plasmid and strain construction

*bcaba1*, *bcaba2*, *bcaba3*, *bcaba4* and *bccpr1* from *B. cinerea* strain ATCC58025 were amplified from plasmids containing codon-optimized sequences for *S. cerevisiae* generated by Otto *et al.* (50). *bmrBM3* (85) and its natural linker were amplified from *BM3* (a kind gift from Dr. Delphine Devriese and Prof. Bart Devreese, Ghent University). *tHMG1* was amplified from the genomic DNA of the Enterol *S. boulardii* strain (28).

Q5 High-Fidelity DNA polymerase (New England Biolabs) was used for PCR of DNA fragments. PCR products were purified using the Wizard® SV Gel and PCR Clean-Up System (Promega). Donor and guide plasmids were generated by Gibson assembly using NEBuilder® HiFi DNA Assembly (New England Biolabs) following the instruction manual provided by the manufacturer. All primers used for plasmid construction are listed in **Table S2**. Plasmids were amplified in *E. coli* cultivated at 37 °C in 5 mL LB medium with ampicillin shaking at 200 rpm. The NucleoSpin Plasmid mini kit (Machery-Nagel) was used for plasmid extraction from *E. coli*.

Genes were genomically introduced by CRISPR/Cas9 mediated gene integration using the LiAC/SS-DNA/PEG protocol as transformation method (86). The Cas9 expression plasmids used in this study was previously described in the report by Cen *et al*. (87) and the gRNA plasmids used are described in **Table S1**. The presence of the introduced genes was verified by colony PCR where DNA was extracted by lysing cells with NaOH (10 mM) for 10 min at 99°C and PCR was performed using the ALLin™ Red Taq Mastermix (highQu). Next, by sub-culturing 3 times in YPD, the Cas9 and guide plasmids were lost. Plasmid loss was verified by spot assay on YPD supplemented with geneticin or nourseothricin. All constructed strains with information about integration sites and verification primers can be found in **Table S1**.

### *S. boulardii* cultivation for ABA measurement

For the comparison of ABA production by the wild-type and engineered *S. boulardii* strains, cultures were grown at 30°C, 33°C or 37°C, while shaking at 200 rpm. A 3 mL preculture in YPD medium was inoculated by picking one yeast colony from a YPD plate. Three to five biological replicates were prepared for each strain. After approximately 18 h of cultivation, main cultures were inoculated at an OD_600_ of 0.1-0.2 and grown for 24 h. Main cultures were prepared in either 50 mL or 20 mL of YPD in 250 mL or 100 mL baffled flasks respectively or in 3 mL of YPD in test tubes or 24-well deep-well plates as specified in the figure legend. After measuring the OD_600_, the cells were harvested by centrifugation (3000xg, 5 min) and the supernatant was collected for ABA measurement (see below).

### ABA measurement in culture supernatant

ABA measurements were performed using a highly sensitive biosensor, engineered by our lab (54). Briefly, this biosensor is a HEK 293T cell line stable transfected with a plasmid containing a PYL1_H87P_ receptor coupled to a VP16 activation domain, the ABA coreceptor ABI1 coupled to a GAL4 binding domain and a plasmid containing a luciferase reporter gene. Presence of ABA brings the VP16 activation domain to the GAL4-dependent promoter, leading to luciferase gene expression. To measure ABA in culture supernatant, reporter cells were seeded at 10.000 cells per well in 96-well plates. One day post seeding, samples were added at a dilution of 1:20 when the medium was refreshed. Positive controls of 8, 4, 2, 1 and 0.5 µM ABA, as well as a negative control were used to create a standard concentration curve. After 24 h, cells were lysed in luciferase lysis buffer (25mM Tris-phosphate pH7.8, 2 mM DTT, 2mM CDTA, 10% glycerol, 1% Triton X-100). To cell lysates, D-luciferin (E1605, Promega) was added and luminescence signals were measured in triplicate using the Glomax® 96 Microplate Luminometer (Promega).

### Quantitative reverse transcription PCR

For gene expression analysis of *bcaba1-4* at different growth temperatures, yeast cells were grown at 30°C for approximately 18 h and main cultures were inoculated at an OD_600_ of 0.1-0.2 and grown for 24 h at either 30°C or 37°C. Subsequently, cells were harvested by centrifugation (Eppendorf centrifuge, 4000 rpm, 2min) and RNA was extracted from cell pellets using the RiboPure™-Yeast Kit (ThermoFisher Scientific) according to the manufacturer’s instructions. RNA concentration and quality were determined with a Nanodrop spectrophotometer (Isogen Life Science). cDNA synthesis was conducted using the SensiFast cDNA Synthesis Kit (GC Biotech) according to the manufacturer’s instructions. qPCR was performed on a Real-Time PCR system (Lightcycler 480, Roche) using the SensiFast SYBR No-Rox Kit containing 1x SYBR green (GC Biotech), 0.3 μM primers (Integrated DNA Technologies), 2 ng cDNA and nuclease-free water. mRNA expression was analyzed using the ΔΔCt method, where expression of *bcaba1*, *bcaba2*, *bcaba3* and *bcaba4* are normalized to expression of *25S rRNA*. Primer sequences are provided in **Table S3**.

### Mice

Male and female wild-type C57BL/6J mice (8-12 weeks old) were obtained from Janvier (France) or bred at Ghent University in specific pathogen–free or germ-free conditions. Mice from Janvier were acclimatized for 2 weeks before initiation of the experiments. The mice were maintained on a 12-h light/dark cycle with free access to water and were fed either a standard chow diet, an ABA-supplemented diet (400 mg/kg, Research diets Inc.) or yeast-supplemented diet (see below), as indicated in the figure legend. All animal procedures were conducted in accordance with the institutional guidelines and approved by the Ethical Committee for Animal Experiments of the VIB site at Ghent University Faculty of Sciences (EC2020-013) or the Ethical Committee for Animal Experiments of the UZ site at Ghent University Faculty of Medicine and Health Sciences (ECD 21-52).

### Yeast-supplemented mouse diet

Yeast-supplemented diet was prepared by mixing standard powdered mouse food (Sniff Bio Services B.V.) with water and yeast pellets creating a porridge-like formulation. In detail, yeast were precultured in 3 mL YPD at 30°C. After approximately 18 h of cultivation, main cultures were prepared in 200 mL of YPD and grown for 24 h at 30°C. The OD_600_ was measured and an equivalent of 15 x 10^9^ CFU were harvested by centrifugation (3000×g, 5 min). The cell pellets were resuspended in 30 mL of drinking water, forming a yeast suspension. This suspension was mixed with 30 grams of powdered mouse diet, resulting in a final concentration of 5 x 10^8^ CFU per gram of food. The mixture was placed in glass jars secured in stainless steel holders within the cages, allowing the mice to consume the diet *ad libitum*. This process was repeated daily, and fresh yeast-supplemented food was given for 1 week before mice were sacrificed and serum was collected. In these experiments, diet supplemented with ABA-producing yeast was compared to diet-supplemented with wild-type yeast or diet supplemented with water, as indicated in the figure legends.

### Administration of yeast to mice via oral gavage

For experiments in which administration of yeast was not performed through the diet, mice received oral gavage with a PBS solution containing the ABA-producing yeast. Conventional mice were gavaged daily with 5 x 10^8^ CFU in 200 μL of PBS for a period of 1 week, while germ-free mice received a single dose of 5 x 10^8^ CFU.

### ABA measurement in serum

ABA measurements were performed using the highly sensitive biosensor, engineered by our lab (54) as described above. However, samples were added at a dilution of 1:10 when the medium was refreshed and positive controls in a lower range were included to create a standard curve (80, 40, 20, 10, 5, 2.5, 1.25 and 0 nM ABA). The standard curve was spiked with ABA-negative serum to correct for sample type-induced effects.

### CFU count in stool

Fecal samples were collected on d 2, 3 and 4 for conventional mice and on d 3, 7 and 13 for germ-free mice. A piece of stool was collected and weighed to determine fecal mass. Fecal samples were then resuspended in 1 mL PBS per 25 mg feces. Fecal suspensions were diluted 1:100 in 1 mL of PBS and 100 µL of these suspensions were spread on solid YPD media containing chloramphenicol (50 mg/L). Plates were incubated at 30°C for 2 d and colonies were counted manually.

### Statistical analysis

Statistical analyses and data visualization were performed using Prism (Graphpad, La Jolla, CA). Data are presented as mean ± standard error of the mean (SEM). The number of biological replicates is indicated by dots in the figure or denoted as “n” in the legend. Data distribution and variance characteristics were considered for statistical testing. For normally distributed datasets, an ordinary one-way ANOVA with correction for multiple testing or a two-tailed unpaired Student’s t-test was used. For non-normally distributed datasets, the Kruskal–Wallis test or Mann–Whitney U test was applied. Statistical significance was defined as p<0.05, with significance levels indicated as * p < 0.05, ** p < 0.01, *** p < 0.001, and **** p < 0.00001. For mouse experiments, no randomization was done, and the investigator was not blinded to the mouse group allocation. The sample size was determined by power analysis using G*power software.

## Material availability

All plasmids and yeast strains described are available upon request.

## Ethics statement

This study was approved by and conducted in accordance with the recommendations of the institutional animal care and use committees at the VIB site of Ghent University Faculty of Sciences or the UZ site of Ghent University Faculty of Medicine and Health Sciences.

## Acknowledgments

We would like to thank all the people of the IRC Animal House Facility, as well as L. Vereecke and his team from the Germ-free Facility at MRB2. We would also like to thank V. Siewers (Chalmers University of Technology, Gothenburg) for her kind supply of *bcaba* plasmids and ABA-producing *S. cerevisiae* strains, and D. Devriese from the lab of B. Devreese for their kind supply of *bmrBM3* DNA (Ugent, Ghent). We would like to thank Charly and Geert from the VIB workshop for their help with the construction of the feeding systems. We would also like to thank all member of the Unit of Molecular Signal Transduction in Inflammation at VIB-UGent for their helpful feedback and suggestions during the preparation of this manuscript.

## Supplementary Figures and Tables

**Figure S1. OD_600_ of strains analyzed for ABA production. (A)** OD_600_ of strains analyzed in Figure 1A, cultivated for 24 h at 30°C in 50 mL of YPD medium. **(B)** OD_600_ of strains analyzed in Figure 2A, cultivated for 24 h at 30°C in 3 mL of YPD medium in 24-well deep well plates. **(C)** OD_600_ of strains analyzed in Figure 2B, cultivated for 24 h at 30°C in 3 mL of YPD medium in test tubes. For more information on genetic composition of strains see **Table 1**.

**Figure S2. Administration of yeast-supplemented diet. (A)** Yeast-supplemented diet was prepared by mixing standard powdered mouse food with water and yeast pellets to form a porridge-like consistency. **(B)** The yeast-supplemented diet was placed into glass jars secured in stainless steel holders to prevent moving or tipping over. **(C)** This feeding system was placed into the cages to allow *ad libitum* consumption.

**Figure S3. ABA measurement in feeding bowls and in yeast colonies retrieved from stool samples of mice fed a yeast-supplemented diet. (A)** Feeding bowls were kept at room temperature for 0, 8 or 24 h. Water was added to the food and following thorough mixing, ABA titers were measured in the supernatant. **(B)** Stool samples from mice fed a yeast-supplemented diet with E-ABA1 (Enterol strain containing *bcaba1-*4) were plated on agar plates and individual yeast colonies were isolated and cultured in 3 mL of liquid medium in test tubes at 30°C for 24 h. ABA titers were measured in the supernatant of these cultures and are presented relative to the ABA titers in the supernatant of E-ABA1 cultures that have not passed through the gut.

**Figure S4. Determination of CFU count in the feces of germ-free mice administered ABA-producing yeast through oral gavage. (A)** Schematic representation showing treatment of the mice. Germ-free mice were given a single oral gavage of 5 x 10^8^ CFU of E-ABA1 on d 0 (yellow dot). The time points for collection of feces are depicted by brown dots. **(B)** Fecal carriage of yeast. Fecal samples were collected as shown in the timelines, plated on YPD media containing chloramphenicol (50 mg/L) and kept at 30°C for 2 d. Data are represented as mean ± SEM (n = 5 mice).

**Figure S5. mRNA expression levels of *bcaba1-4* in TABA3 strain. (A)** mRNA expression of *bcaba1, bcaba2, bcaba3* and *bcaba4* normalized to expression of *25S rRNA*, measured in the cell pellets of a 24 h culture of TABA3, grown at 30°C in 3 mL of YPD in test tubes. TABA3 is a *S. cerevisiae* strain producing ABA. More information on the genetic composition can be found in **Table 1**. Average expression was calculated from 3 independent biological replicates. Data are represented as mean ± SEM and statistically analyzed using a Student’s t-test. * p < 0.05, ** p < 0.01, *** p < 0.001, **** p < 0.0001.

**Figure S6. mRNA expression levels of *bcaba1* and *bcaba3* and OD_600_ of strains E-ABA1, E-ABA2, E-ABA3, E-ABA4 and E-ABA5.** Strains were grown at 30°C or 37°C in 3 mL YPD in 24-well deep well plates. **(A-B)** mRNA expression of *bcaba1* and *bcaba3* were normalized to expression of *25S rRNA*, measured in the cell pellets of a 24 h culture. Average expression was calculated from 3 independent biological replicates. Data are represented as mean ± SEM. **(C)** OD_600_ of strains analyzed in **Figure S5A,B** and Figure 6A**,B**.

**Table S1. Details about genomic integrations**

**Table S2. Primers for plasmid construction and amplification of donor DNA**

**Table S3. qPCR primers**

