## Supplementary figures and images for "Multi-Step Pathway Engineering in Probiotic *Saccharomyces boulardii* for Abscisic Acid Production in the Gut"

### Fig. S1

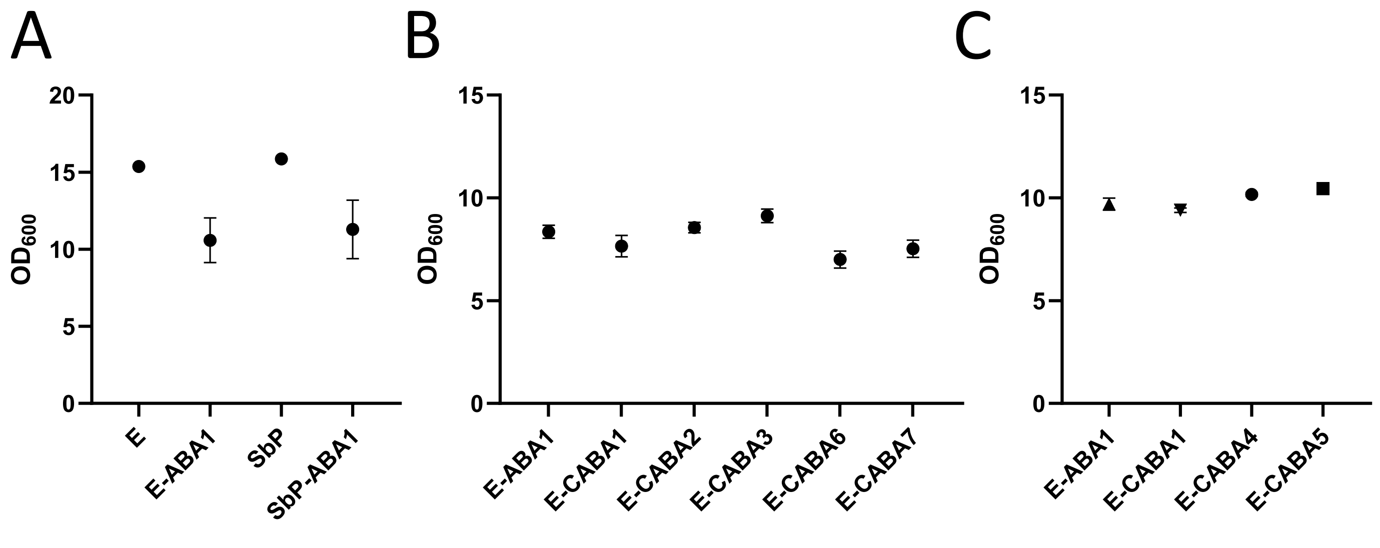

### Fig. S2

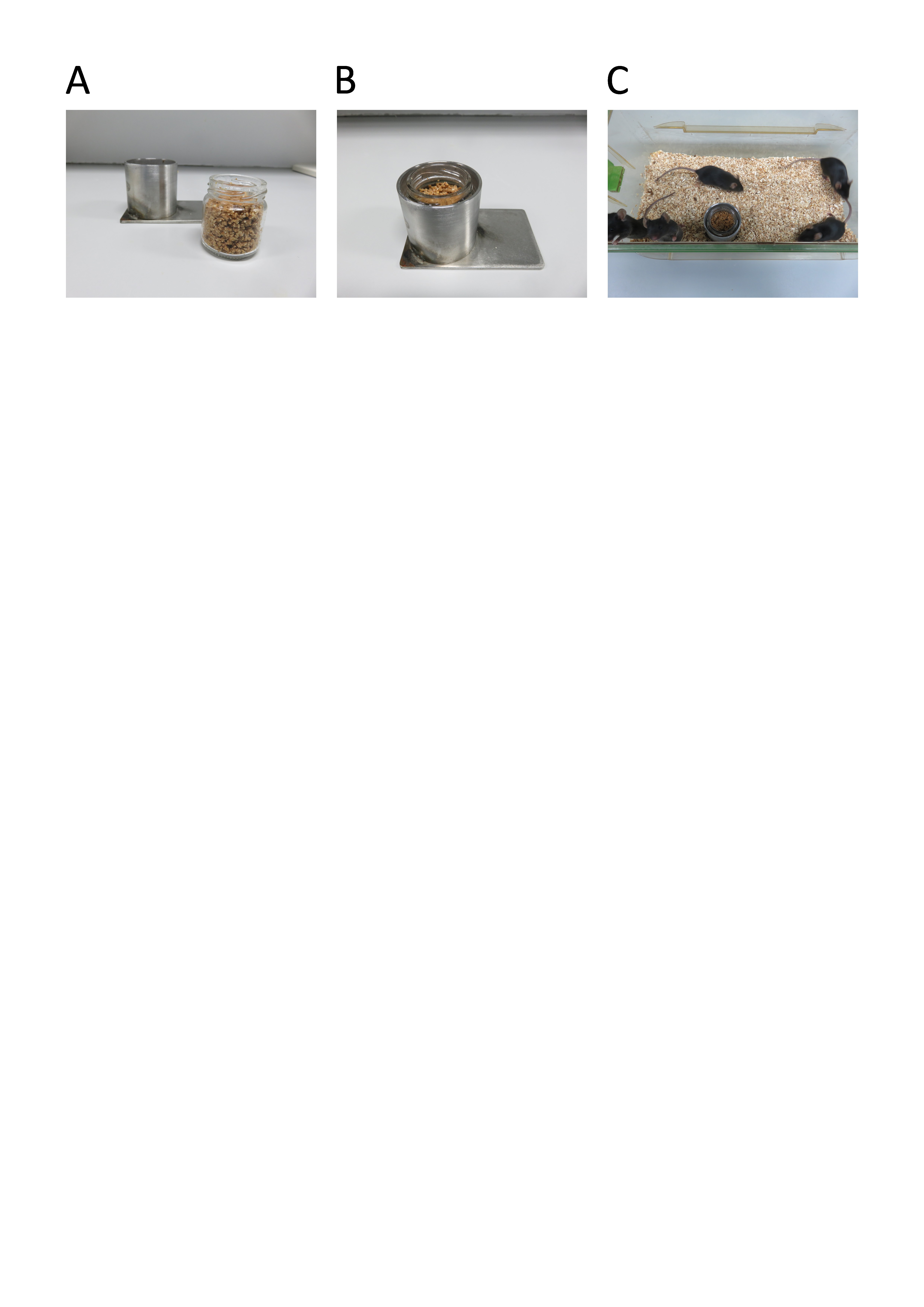

### Fig. S3

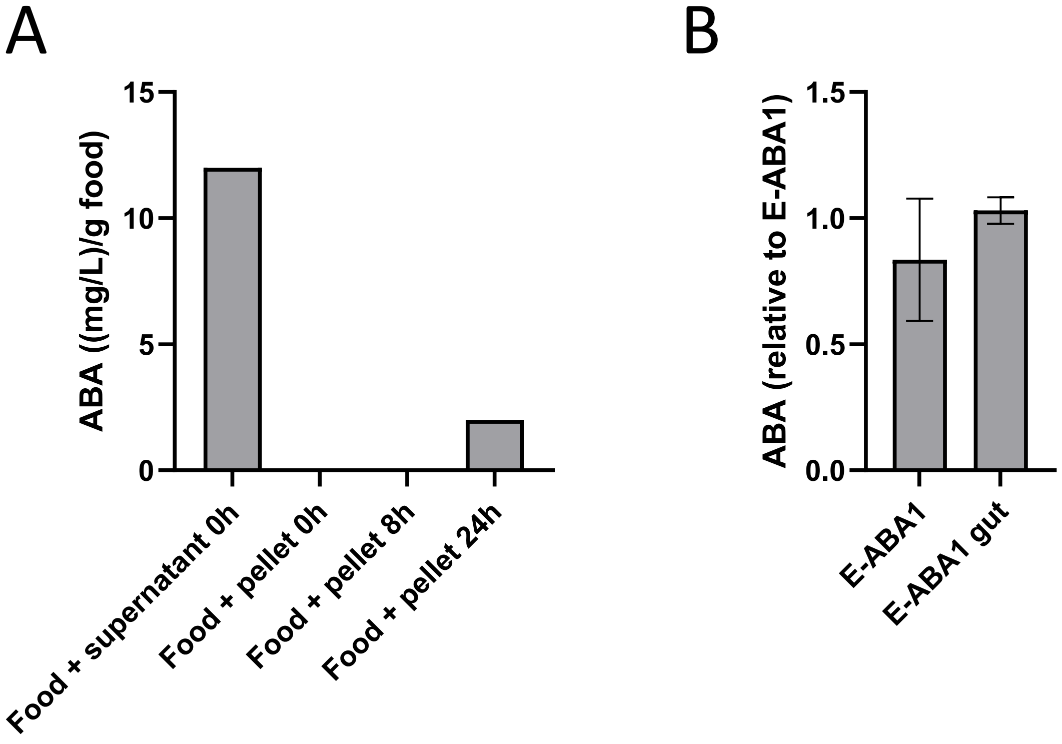

### Fig. S4

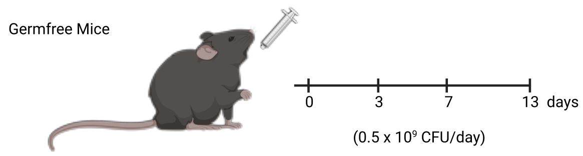

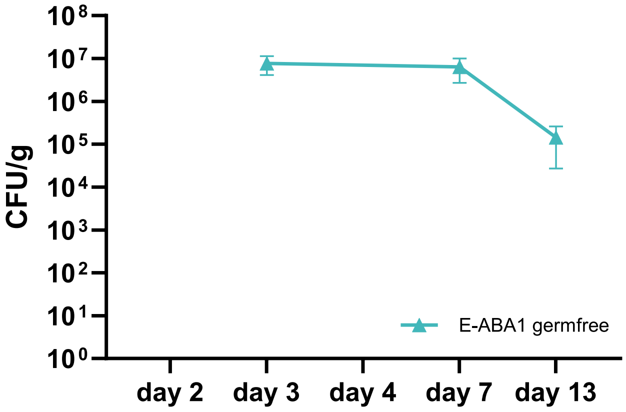

### Fig. S5

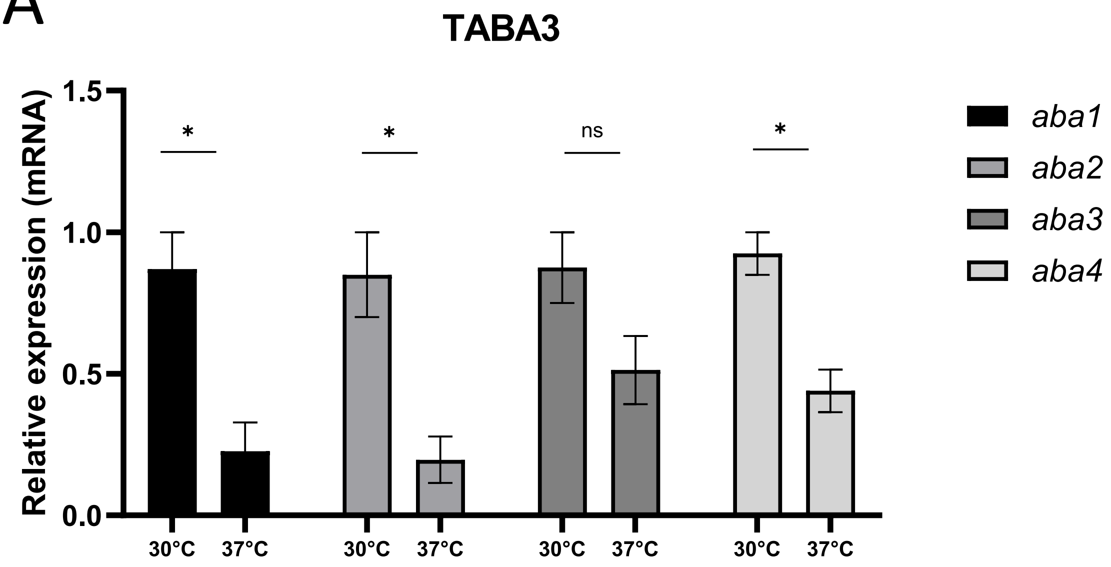

### Fig. S6

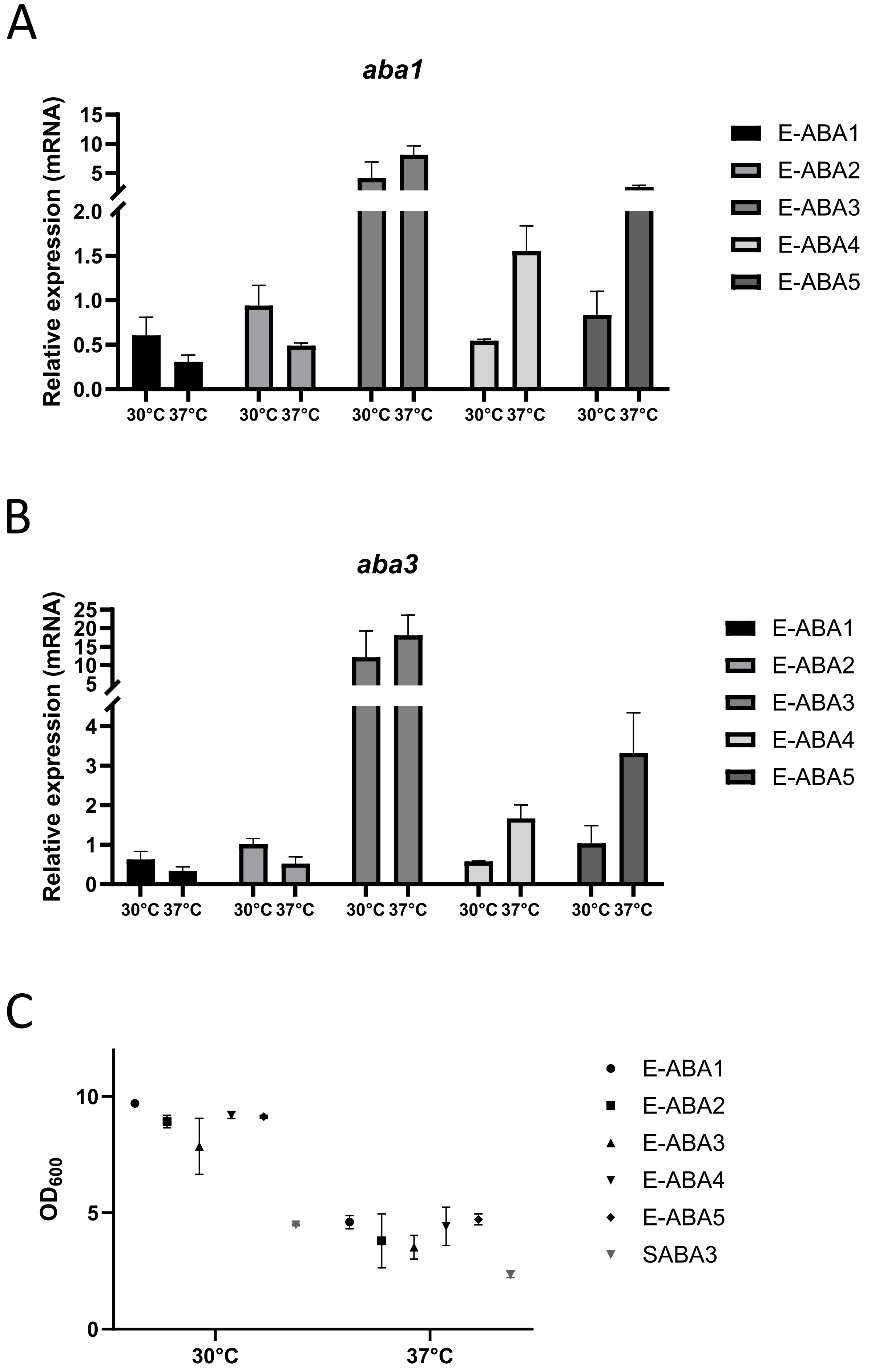
