## Supplementary material for "Multi-Step Pathway Engineering in Probiotic *Saccharomyces boulardii* for Abscisic Acid Production in the Gut": Table S1

**Table S1. Details about genomic integrations**

| **Yeast strain** | **Yeast backbone** | **Integrated expression casettes** | **Integration site (gRNA sequence) (Reference)** | **Verification primers** |
| --- | --- | --- | --- | --- |
| E-ABA1 | E | P*_TDH3_*-*bcaba1*-*2A*-*bcaba3*-T*_ENO2_* P*_TEF1_*-*bcaba2*-*2A*-*bcaba4*-T*_CYC1_* | IV-1 (CAAAAGCGACACGTCGTCTG) (1) | F: GCCTGGCATTGGTGAAAAGGC R: ACAGGTTTCGTACTCTCTTGC |
| E-ABA2/3/4/5 | E-ABA1 | P*_TDH3/TEF1/RPL18B/SAC6_*-*bcaba1-2A‑bcaba3-*T*_ADH1_* | XII-1 (GAGTTTCATAACGCGTTACA) (2) | F: ATCCCATATGTGACGCAGCG R: GTAAGAGAGGTTAATGTCCTC |
| E-CABA1 | E-ABA1 | P*_TEF1_*-*cpr1*-T*_CYC1_* | II-1 (ATCAACCACAGTGAACGCCG) (1) | F: TCGAATCCAGAATCAGATACC R: ACATTGCTATCGCCTCACTGC |
| E-CABA2 | E-ABA1 | *cpr1* | aba1 (TCTGAAGGTACAGAATACAA) (This study) | F: TGTGTTGGTAAAACAGTTGC R: GCAGTTGCAGCTTTAATTGG |
| E-CABA3 | E-ABA1 | *cpr1* | aba2 (AAATCATAATTAGATTTTCT) (This study) | F: GGGTCCAAGAGGTTGTATCG R: TATCCTCTTGGATATCTGC |
| E-CABA4 | E-ABA1 | *bmr* | aba1 (TCTGAAGGTACAGAATACAA) (This study) | F: TGTGTTGGTAAAACAGTTGC R: GCAGTTGCAGCTTTAATTGG |
| E-CABA5 | E-ABA1 | *bmr* | aba2 (AAATCATAATTAGATTTTCT) (This study) | F: GGGTCCAAGAGGTTGTATCG R: TATCCTCTTGGATATCTGC |
| E-CABA6 | E-CABA4 | *cpr1* | aba2 (AAATCATAATTAGATTTTCT) (This study) | F: GGGTCCAAGAGGTTGTATCG R: TATCCTCTTGGATATCTGC |
| E-CABA7 | E-CABA5 | *cpr1* | aba1 (TCTGAAGGTACAGAATACAA) (This study) | F: TGTGTTGGTAAAACAGTTGC R: GCAGTTGCAGCTTTAATTGG |
| E-TABA2/3/4/5 | E-ABA2/3/4/5 | P*_TEF1_*-*tHMG1*-T*_CYC1_* | XVI-1(GTTAACTCTGAATGTATGTG) (This study) | F: CCTAAACAACGACCCGGAAATA R: CATGCGCGGCAATTTGATAG |
| E-TCABA2/3 | E-TABA2/3 | P*_TEF1_*-*cpr1*-T*_CYC1_* | II-1 (ATCAACCACAGTGAACGCCG) (1) | F: TCGAATCCAGAATCAGATACC R: ACATTGCTATCGCCTCACTGC |
