## Supplementary material for "Multi-Step Pathway Engineering in Probiotic *Saccharomyces boulardii* for Abscisic Acid Production in the Gut": Table S2

**Table S2. Primers for plasmid construction and amplification of donor DNA**

| **Constructed plasmid** | **Template (reference)** | **Primer** | **Sequence (upper-case letters represent part binding template; lower-case letters represent overhang)** |
| --- | --- | --- | --- |
| p426-*aba1-aba2-aba3-aba4* |  | *bcaba1-4* Fwd | tttgggcaagtggggttgagtggagtgtcaAAGTTTATCATTATCAATACTCG |
|  |  | *bcaba1-4* Rev | gcacgttcttatgtccgtaaaaacggatataTTCGAGCGTCCCAAAACC |
|  | p416-*aba1+2* (1) | *bcaba1* Fwd | tcctgataggactctgtaagtgttaAGTTTATCATTATCAATACTCG |
|  |  | *bcaba1* Rev | caccagccaatttcaacaaagaaaaattagtagcaccTTTGTATTCTGTACCTTCAG |
|  | p416-*aba1+2* (1) | *bcaba2* Fwd | GGATCCAGTGCTTTTAACTAAG |
|  |  | *bcaba2* Rev | tcaccacataattttaataaatcataattagatttTCTTGGAACTTCTTTTAACATAAC |
|  | pCfB3035-*aba3-aba5* (1) | *bcaba3* Fwd | ttttctttgttgaaattggctggtgatgttgaattgaatccaggtccaATGCAACAAGTTATTACTCAAAC |
|  |  | *bcaba3* Rev | attcttagttaaaagcactggatccTTAAACTGGAACTTCAAAATG |
|  | pCfB2909-*aba2-aba4* (1) | *bcaba4* Fwd | atgatttattaaaattatgtggtgatgttgaatctaatccaggtccaATGTCTTCACAACCTTTTAC |
|  |  | *bcaba4* Rev | ggaaactcgtcatattctaccaaggCTTCGAGCGTCCCAAAACCTTC |
|  | p426 backbone (2) |  |  |
| pCR2.1-*TDH3*p-*aba1-aba3*-*ADH1*t |  | *TDH3-bcaba1-3* Fwd | acgggcaatcagaatctgtaacaagcgccatttttttttctgtatcgggccctccttactgctctccttcatCAAGCTTGGTACCGAGC |
|  |  | *TDH3-bcaba1-3* Rev | ccaagtggcaaaagcgttagacgcagtacaaggacgcgttaagaaaaatttcgagagagtcgccgatagCTCACTATAGGGCGAATTGG |
| pCR2.1-*TEF1*p-*aba1-aba3*-*ADH1*t |  | *TEF1-bcaba1-3* Fwd | acgggcaatcagaatctgtaacaagcgccatttttttttctgtatcgggccctccttactgctctccttcatCAAGCTTGGTACCGAGC |
|  |  | *TEF1-bcaba1-3* Rev | ccaagtggcaaaagcgttagacgcagtacaaggacgcgttaagaaaaatttcgagagagtcgccgatagCTCACTATAGGGCGAATTGG |
| pCR2.1-*RPL18B*p-*aba1-aba3*-*ADH1*t |  | *RPL18B-bcaba1-3* Fwd | acgggcaatcagaatctgtaacaagcgccatttttttttctgtatcgggccctccttactgctctccttcatCAAGCTTGGTACCGAGC |
|  |  | *RPL18B-bcaba1-3* Rev | ccaagtggcaaaagcgttagacgcagtacaaggacgcgttaagaaaaatttcgagagagtcgccgatagCTCACTATAGGGCGAATTGG |
| pCR2.1-*SAC6*p-*aba1-aba3*-*ADH1*t |  | *SAC6-bcaba1-3* Fwd | acgggcaatcagaatctgtaacaagcgccatttttttttctgtatcgggccctccttactgctctccttcatCAAGCTTGGTACCGAGC |
|  |  | *SAC6-bcaba1-3* Rev | ccaagtggcaaaagcgttagacgcagtacaaggacgcgttaagaaaaatttcgagagagtcgccgatagCTCACTATAGGGCGAATTGG |
|  | p426-*aba1-aba2-aba3-aba4* (This study) | *TDH3-bcaba1* Fwd | aacacacataaacaaacAAAATGTCTAACTCAATCTTGAAT |
|  |  | *TEF1-bcaba1* Fwd | atctaagttttaattacAAAATGTCTAACTCAATCTTGAAT |
|  |  | *RPL18B-bcaba1* Fwd | agaagaaaacaaaaaacAAAATGTCTAACTCAATCTTGAAT |
|  |  | *SAC6-bcaba1* Fwd | aaggagtacaccaaaacacAAAATGTCTAACTCAATCTTGAAT |
|  |  | *ADH1-bcaba3* Rev | ataaatcataagaaattcgCTTAAACTGGAACTTCAAAATGTGTC |
|  | pCR2.1-TDH3p-NotI-XhoI-ADH1t (3) |  |  |
|  | pCR2.1-TEF1p-NotI-XhoI-ADH1t (3) |  |  |
|  | pCR2.1-RPL18Bp-NotI-XhoI-ADH1t (3) |  |  |
|  | pCR2.1-SAC6p-NotI-XhoI-ADH1t (3) |  |  |
|  | pCfB2904-ABA1-CPR1 (1) | *bccpr1* Fwd | aagttttattgcagcatcctttaatggcaaagatggcgcgataaggacgaaaccggcaaatcccgagcGCCGCACACACCATAGCTTC |
|  |  | *bccpr1* Rev | agccatgccacaaagcggtatgtgccattaatttcttcttttctcccctttcgcctttgagtttct CTTCGAGCGTCCCAAAACCTTC |
|  | pCfB2904-ABA1-CPR1 (1) | *bcaba1-bccpr1* Fwd | cttggaatcaaacatgttggttaaggttactgaatctgaaggtacagaatacaaaACAAAGGGTAAATACTGGGGTG |
|  |  | *bcaba1-bccpr1* Rev | ctggattcaattcaacatcaccagccaatttcaacaaagaaaaattagtagcaccAGACCAAACATCTTCTTGGTATTG |
|  | pCfB2904-ABA1-CPR1 (1) | *bcaba2-bccpr1* Fwd | ttaccaaacatggaataagccagatatgtgggttatgttaaaagaagttccaagaACAAAGGGTAAATACTGGGGTG |
|  |  | *bcaba2-bccpr1* Rev | gacctggattagattcaacatcaccacataattttaataaatcataattagatttAGACCAAACATCTTCTTGGTATTG |
|  | BMR (gift from Dr. Devriese and Prof. Devreese, Ugent) | *bcaba1-bmr* Fwd | cttggaatcaaacatgttggttaaggttactgaatctgaaggtacagaatacaaaATTCCTTCACCTAGCACTGAACAG |
|  |  | *bcaba1-bmr* Rev | ctggattcaattcaacatcaccagccaatttcaacaaagaaaaattagtagcaccCCCAGCCCACACGTCTTTTGC |
|  | BMR (gift from Dr. Devriese and Prof. Devreese, Ugent) | *bcaba2-bmr* Fwd | ctggattcaattcaacatcaccagccaatttcaacaaagaaaaattagtagcaccCCCAGCCCACACGTCTTTTGC |
|  |  | *bcaba2-bmr* Rev | gacctggattagattcaacatcaccacataattttaataaatcataattagatttCCCAGCCCACACGTCTTTTGC |
| IS16.1-*tHMG1* |  | IS16.1-*tHMG1* Fwd | CTTATTTCTTCCTTGCGCACTATTT |
|  |  | IS16.1-*tHMG1* Rev | GCGTAATTGAGCTTGGCTTTC |
|  | genomic DNA Enterol (4) | *tHMG1* Fwd | taagttttctagaactagtggatccATGGTTTTAACCAATAAAACAGTC |
|  |  | *tHMG1* Rev | tgacataactaattacatgactcgaTTAGGATTTAATGCAGGTGAC |
|  | p426 backbone (2) |  |  |
