## Supplementary material for "Multi-Step Pathway Engineering in Probiotic *Saccharomyces boulardii* for Abscisic Acid Production in the Gut": Table S3

**Table S3. qPCR primers**

| **Gene** | **Primer** | **Sequence** |
| --- | --- | --- |
| *aba1* | *aba1* Fwd | AGGTGACTGGTATTTGACTGG |
|  | *aba1* Rev | CGATCAAAGTTTGCAATTGAGC |
| *aba2* | *aba2* Fwd | CTGAAATCCCAGGTTCATGGG |
|  | *aba2* Rev | CATCCCATGCTCTGTCACC |
| *aba3* | *aba3* Fwd | AAATCTTCGCTTGTGCATGGG |
|  | *aba3* Rev | GCATCCAAATCGTAACCCAAC |
| *aba4* | *aba4* Fwd | CGGTGAAGTTGAAGCATGG |
|  | *aba4* Rev | GCAGATATCCAAGAGGATA |
| 25S rRNA | 25S Fwd | AAGAGTGCGGTTCTTTG |
|  | 25S Rev | TACTTACCGAGGCAAGCTACA |
